# Reduced mitochondrial transcription sensitizes acute myeloid leukemia cells to BCL-2 inhibition

**DOI:** 10.1101/2022.09.23.509145

**Authors:** Laleh S. Arabanian, Jenni Adamsson, Anke Unger, Raffaella Di Lucrezia, Tim Bergbrede, Arghavan Ashouri, Erik Larsson, Peter Nussbaumer, Bert M. Klebl, Lars Palmqvist, Claes M. Gustafsson

## Abstract

Overcoming drug-resistance and the subsequent relapse that often occurs with monotherapy is crucial in the treatment of acute myeloid leukemia. We here demonstrate that therapy-resistant leukemia initiating cells can be targeted using a novel inhibitor of mitochondrial transcription (IMT). The compound inhibits mitochondrial RNA polymerase activity and sensitizes the resistant population to the induction of apoptosis. *In vitro* studies on acute myeloid leukemia cells demonstrate that IMT prevents cell proliferation, and together with a selective BCL-2 inhibitor, venetoclax, induces apoptosis and suppress oxidative phosphorylation (OXPHOS) synergistically. AML mouse models treated with IMT in combination with venetoclax show prolonged survival in venetoclax-resistant models. Our findings suggest that certain therapy-resistant leukemia cell populations display a unique dependency on mitochondrial transcription and can be targeted with IMT.

## Introduction

Acute myeloid leukemia (AML) comprises a devastating and heterogeneous group of diseases, for which patient survival rates are dismal despite several new treatment options. Current cares are still mainly based on unselective chemotherapy but for most patients also with addition of more directed treatments after patient stratification based on genetic aberrations found at diagnosis. However, these strategies lead to cures in only a fraction of AML patients due to therapy resistance developed by the leukemic cells leading to relapse. Curative treatment usually needs allogenic stem cell transplantation, which is not applicable in many cases, highlighting the critical need to develop different approaches. The anti-apoptotic molecule BCL-2 has been a therapeutic target in lymphoid and myeloid leukemia, as in many cancers overexpression of BCL-2 results in dysregulation of apoptosis (1). ABT-199 (venetoclax), is a specific BH3-mimetic that causes apoptosis by preventing BCL-2 from binding pro-apoptotic BAX and BAK1 proteins (2). However, complete responses occur in just ∼20% of AML patients treated with Venetoclax (3). In addition, many patients relapse, with regrowth initiated by therapy-resistant leukemic clones, suggesting that single agent administration is unlikely to yield lasting responses in most cases. Venetoclax is therefore used in combination with conventional therapies. Together with hypomethylating agents, Venetoclax is now standard of care for patients deemed unfit for intensive induction therapy (4).

Resistance to venetoclax may result from overexpression of the anti-apoptotic proteins MCL1 or BCL2L1 (5), or through changes in levels of other anti-apoptotic molecules (6). Aberrant pro-survival regulation can also be achieved by inactivation of TP53 function, which controls many aspects of apoptosis (6, 7). MCL1-dependent resistance to venetoclax can be overcome when the compound is combined with the MDM2 inhibitor RG7388, which causes p53 activation. A combination of BCL-2 inhibition and TP53 activation may thus provide an interesting therapeutic approach for treatment of AML (8).

Other combination approaches that promote apoptosis may also be of interest. The majority of ATP required for leukemia stem cell (LSC) function is generated by oxidative phosphorylation (OXPHOS) (9, 10). Interestingly, venetoclax-resistant leukemic cells display increased levels of OXPHOS and are protected from mitochondrial stress (7). These data suggest approaches to specifically target OXPHOS in leukemia may be an attractive option. In support of this notion, inhibition of mitochondrial protein translation by antibiotics that reduce OXPHOS capacity results in a selective cytotoxic effect on leukemic cell when used in combination with tyrosine kinase inhibitors (11). Furthermore, LSCs from *de novo* AML patients display a unique dependency on amino acid metabolism to fuel OXPHOS. When venetoclax is combined with the hypomethylating agent azacytidine, the two compounds act in concert to inhibit amino acid uptake, leading to a drop in OXPHOS and thus cell death (12). More recently, AML therapy using a combination of cytarabine (AraC) and venetoclax has highlighted the central role of mitochondrial adaptation during treatment (10, 13), indicating the need for a mitochondrial directed combination therapy for AML. Such a therapy may help to overcome existing treatment resistance.

Recently we developed novel, first-in-class inhibitors of mitochondrial transcription (IMTs) (14). IMT treatment specifically targets mitochondrial RNA polymerase (POLRMT) and reduce transcription of mitochondrial DNA, which codes for thirteen key components of the OXPHOS machinery, including subunits of respiratory complexes I, III, IV, and the ATP synthase. Consequently, IMT treatment impairs OXPHOS system biogenesis in a dose-dependent manner and decreases cell viability in various tumor cells. Surprisingly, several weeks of per oral IMT treatment is well tolerated in mice and does not cause toxicity in normal tissues, despite inducing a strong anti-tumor response in human cancer xenografts (14).

Here we examine AML cell lines for their sensitivity for treatment with IMT. We find that a combination of IMT and venetoclax displays a synergistic on cell growth and induction of apoptosis. We demonstrated the efficacy of this approach in leukemia mouse models generated by human AML cell lines as well as primary samples from AML patients.

## Results

### Synergistic effect of IMT in combination with venetoclax on AML cell lines

We were curious whether our inhibitors of <u>m</u>itochondrial transcription (IMTs) (14) could be effective against AML cells. The coumarin class of IMTs was optimized by medicinal chemistry to generate lead IMT compounds with improved metabolic stability in mice, such as LDC204857, herein simply referred to as IMT (Fig 1A, Fig. S1 and Table S1). IMT is a close analog of previously described IMT1B (Fig. S1) with a highly similar biochemical and cellular inhibition profile (14). It shows an extended half-life *in vivo* due to its increased metabolic stability (Table S1). Thus, by using IMT we could consistently reduce the mouse dose to 50 mg/kg per oral once daily, which is very well tolerated and does not exert any adverse effects in mice.

**Figure 1.**
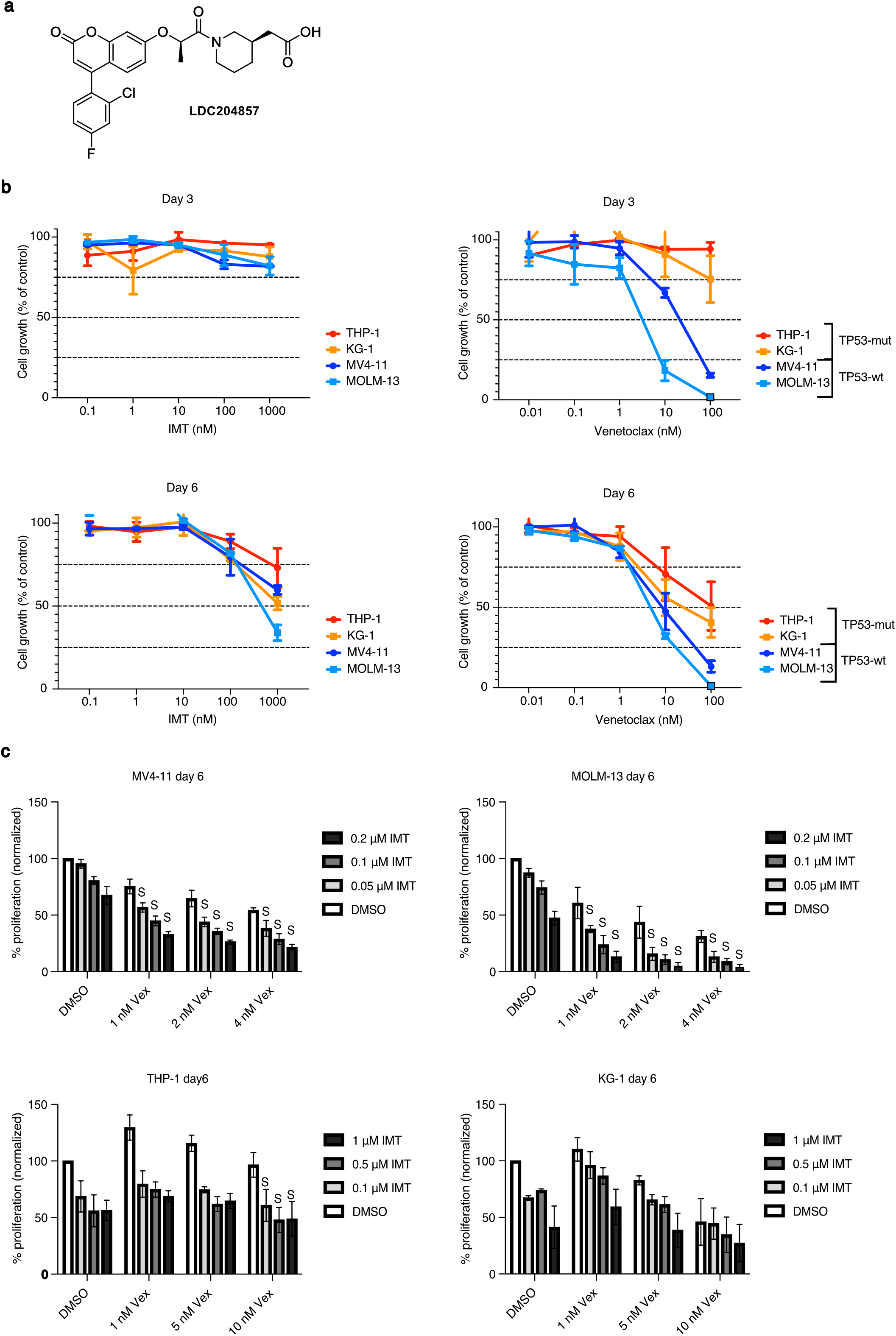
IMT+venetoclax combination treatment synergistically inhibits cell viability/growth of AML cell lines *in vitro*. (**A**) Chemical structure of IMT (LDC204857) (**B**) AML cell lines were treated with single agents IMT or venetoclax at a range of concentrations (0.1–1000 nM and 0.01–100 nM, respectively), and assessed by cell viability tests after three and six days. (**C**) Synergism analysis of AML cell lines treated with a combination of IMT (0.05, 0.1, 0.2 μM for MV4-11 and MOLM-13, and 0.1, 0.5, and 1 μM for THP-1 and KG-1) and venetoclax (Vex; 1, 2, 4 nM for MV4-11 and MOLM-13, and 1, 5, 10 nM for THP-1 and KG-1) for six days. ‘S’ represents synergy of the combination treatment, evaluated using the Bliss independent model. Proliferation percentages are normalized to DMSO treated. All results are from three independent experiments.

Using four AML cell lines with different expression of TP53 (MV4-11 and MOLM-13 expressing wild type TP53; THP-1 and KG-1 expressing mutated TP53), we monitored the effects of increasing concentrations of IMT on cellular growth using a luminescence-based cell viability assay. Only marginal effects were observed after a 72-h period of exposure to IMT, but all cell lines displayed dose-dependent antileukemic activity after 144 h of treatment (Fig. 1B), in line with our previous observations (14). We also monitored venetoclax sensitivity by assessing cell viability, and noted that, in line with published work (15), a dose-dependent reduction in cell growth occurs after three days and becomes more pronounced after 6 days (Fig. 1B).

To then investigate whether IMTs show a synergistic effect when combined with venetoclax, we conducted a combination study in AML cell lines (Fig. 1C). After six days of treatment, we found a clear synergistic effect on cell proliferation in MV4-11 and MOLM-13 cell lines, which express wild type TP53, as determined by combination analysis (Bliss-independence model (16)). Treatment of THP-1 and KG-1 cell lines, which harbor mutated TP53, does not show a synergistic effect even when using higher concentrations of both IMT and venetoclax (Fig. 1C, additional treatment data are shown in Fig. S2, Bliss value calculations for both time points are summarized Fig. 1 - Source Data 1.

### Combination of IMT and venetoclax synergistically induces apoptosis and loss of mitochondrial membrane potential in MV4-11 and MOLM-13 cells

To investigate the factors involved in the inhibition of cell growth with IMT and venetoclax, AML cell lines were treated with single agents (IMT or venetoclax) or a combination of the two (Fig. 2A), and the stages of apoptosis were monitored using Annexin V and PI staining. No detectable increase in apoptosis was induced by IMT exposure alone in any AML cell line. Venetoclax treatment resulted in both early and late apoptosis induction in MV4-11 and MOLM-13 cell lines, and employing a combination of IMT and venetoclax significantly enhanced apoptosis induction in these wild-type cell lines. On the other hand, combination treatment did not induce apoptosis in THP-1 and KG-1 mutated cell lines (Fig. 2A).

**Figure 2.**
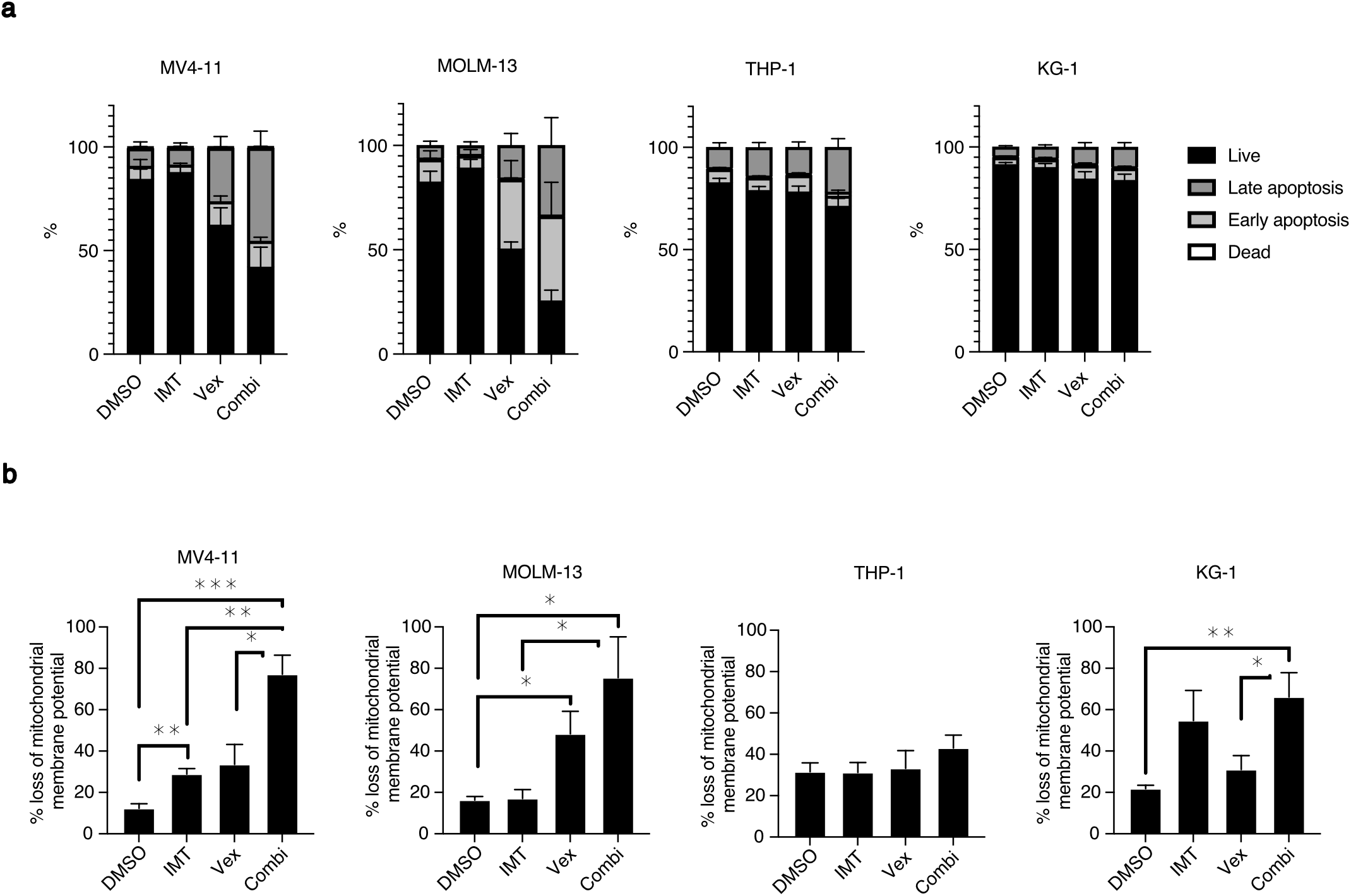
Combination treatment significantly enhances apoptosis induction in AML cell lines. AML cell lines were treated with IMT (500 nM) and venetoclax (5 nM) for 72–96 h. (**A**) Apoptosis was determined by Annexin V/PI staining of the cells and analyzed by flow cytometry. (**B**) Loss of mitochondrial membrane potential was calculated as the percentage of depolarized (apoptotic) cells out of the total cell population. Error bars represent standard error of the mean where * represents P < 0.05, **P < 0.01, and ***p < 0.001.

One distinctive feature of apoptosis is the disruption of mitochondrial function, which can be measured as loss of mitochondrial membrane potential due to membrane depolarization. As a combination of IMT and venetoclax could synergistically induce apoptosis, we were curious whether combination drug treatment thus affects mitochondrial membrane potential. We observed that venetoclax-sensitive cell lines (MV4-11 and MOLM-13) showed a dramatic loss of mitochondrial membrane potential when treated with both venetoclax and IMT (Fig. 2B). On the other hand, mitochondrial membrane potential in THP-1 cells was not affected by single nor by combination treatment. Interestingly, although KG-1 was venetoclax insensitive, the combination treatment significantly enhanced loss of mitochondrial membrane potential (Fig. 2B). These data suggest that a combination of IMT and venetoclax is cytotoxic and synergistically induces cell death in sensitive cells, presumably through disruption of mitochondrial function.

### Combination of IMT and venetoclax significantly suppresses OXPHOS in AML cells

We next studied whether combination treatment with IMT and venetoclax affects OXPHOS in AML cells. OXPHOS was evaluated by determining oxygen consumption rate (OCR) (Fig. 3A), which was measured first as basal respiration and then oligomycin (OMY)-induced leak respiration (ATP-linked respiration) after the addition of oligomycin to inhibit ATP synthase. Stepwise carbonyl cyanide m-chlorophenyl hydrazone (CCCP) titrations were subsequently carried out to reach the electron transport chain respiratory capacity state. Finally, antimycin a (AMA) was added to inhibit complex I/III of the electron transport chain and to terminate respiration, allowing for the determination and correction of residual oxygen consumption, which is indicative of non-mitochondrial oxygen consumption (Fig. 3A). We noted a tendency to inhibition of OXPHOS respiration in venetoclax-sensitive MOLM-13 and MV4-11 cells when combination treatment was used (Fig. 3B). In comparison, OXPHOS levels in venetoclax-resistant THP-1 cells were unaffected in the presence of single compound or combination treatments, confirming that the inhibition we observed is not a general effect of drug treatment (Fig. 3B). In addition, there was no conclusive OCR data obtained for KG-1 cell line.

**Figure 3.**
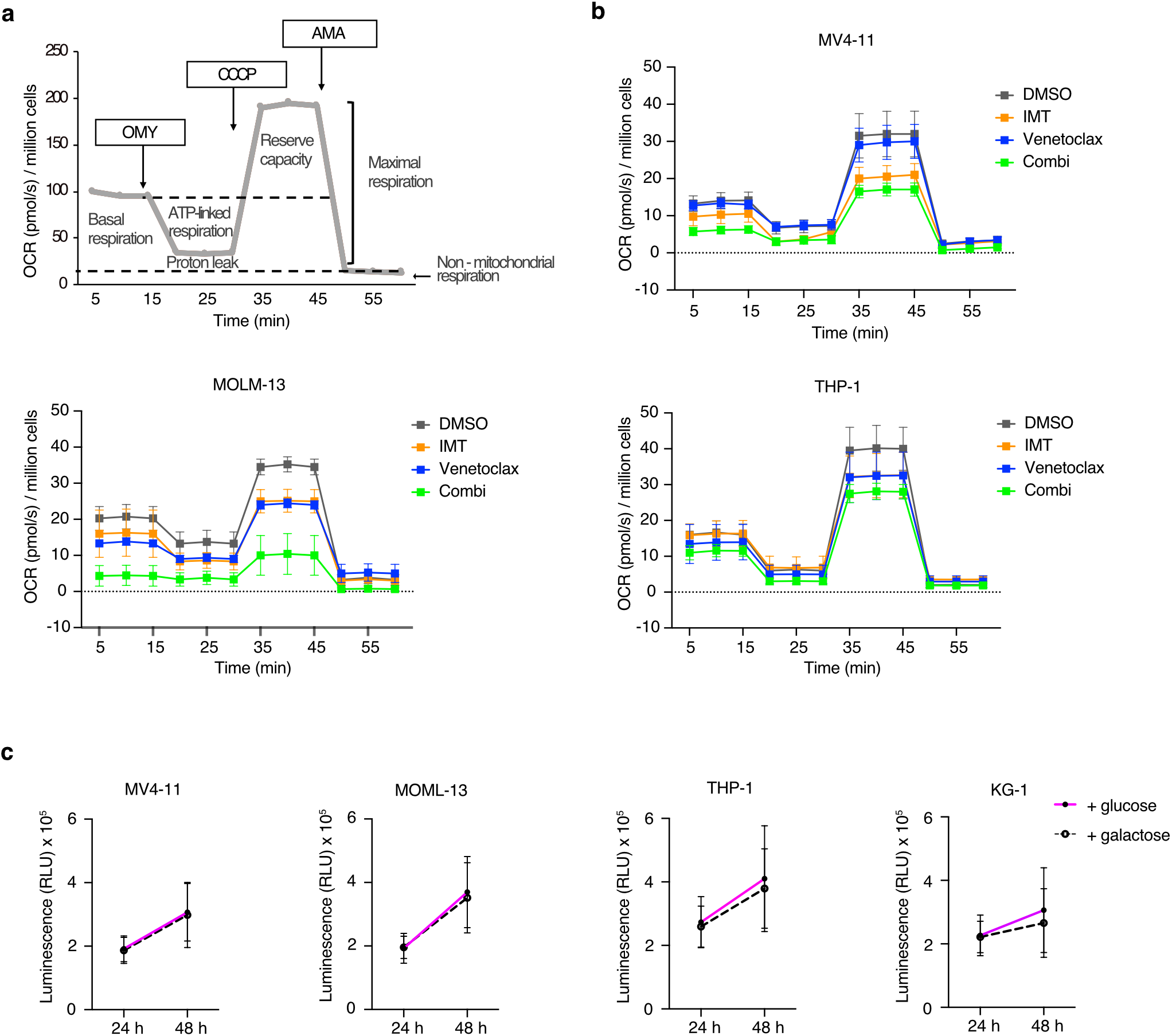
Combination of IMT and venetoclax dramatically decreases OXPHOS (in a glycolysis-independent manner) in sensitive cell lines. (**A**) Representative graph of an OXPHOS measurement using mitostress compounds: oligomycin (OMY), carbonyl cyanide m-chlorophenyl hydrazone (CCCP), and antimycin (AMA). (**B**) AML cell lines (MV4-11, MOLM-13, and THP-1) were treated with either IMT (500 nM), venetoclax (5 nM), or a combination of both for four days. DMSO was added to control wells. Oxygen consumption rate (OCR) was measured with an Oroboros O2K instrument at physiological glucose levels (5 mM) using mitostress compounds. (**C**) AML cell lines were cultured in growth medium supplemented with either 5 mM glucose or galactose, and cell viability tests (ATP measurements) were performed after 24 and 48 h. All results are from at least three independent experiments. Error bars show ± SEM. Statistical tests on all comparisons in (C) showed non-significant differences.

In normal hematopoietic cells, glycolysis is upregulated when OXPHOS is blocked (17). In contrast to normal hematopoietic cells, high energy turnover AML cells rely predominantly on OXPHOS due to poorly regulated metabolic adaptation. To verify the reliance on OXPHOS, AML cells were grown in growth medium supplemented with either glucose or galactose (Fig. 3C). Two ATP molecules are produced when cells use glucose for glycolysis, but if galactose is used instead, no ATP is generated (18). We observed no significant decrease in the level of ATP production (Fig. 3C) nor in cell counts (Fig. S3A) in the absence of glucose, indicating that glycolysis is not a major source of energy in AML cells. To test whether AML cells use glycolysis under treatment conditions, OCR was measured in the presence or absence of a high concentration of glucose (Fig. S3B). OCR measured for basal respiration was not significantly different with excess amounts of glucose, again indicating the inability of AML cells to switch to glycolysis when OXPHOS is disrupted.

To assess whether AML cells could rely on glycolysis for ATP generation when OxPhos was inhibited by IMT, we cultured AML cells in medium with galactose instead of glucose, a condition that limits glycolysis-dependent ATP production. Even in these glycolytic conditions, we observed minimal ATP production and cell viability, suggesting that AML cells are poorly adapted to switch to glycolysis when mitochondrial function is impaired. This finding underscores the high dependence of AML cells on OxPhos for energy production and supports the proposed mechanism of IMT efficacy, as it appears to exploit the inability of AML cells to compensate through glycolysis when mitochondrial transcription is inhibited.

### Combination of IMT and venetoclax synergistically delays leukemia initiation in cell-derived leukemia mouse models

Following the confirmed synergy between IMT and venetoclax in AML cell lines, we proceeded to test the dual inhibition of mitochondrial transcription and BCL-2 in an AML cell-derived xenograft (CDX) transplantation mouse model. First, the pharmacokinetic profile of IMT was determined in plasma, liver, and bone marrow (BM) of mice, 1, 8, and 24 h post-administration (Fig. S4A). As described above (Fig. S1), the IMT mouse PK profile was superior to IMT1B. IMT was detected in all tissues at all time points when administered alone. To determine the concentration of IMT and venetoclax when administered in combination, plasma samples were then collected 8–11 h post-administration of single or combination formulations and the concentrations of IMT and/or venetoclax in samples were measured (Fig. S4B). Values obtained were all within the effective range, as combination treatment did not affect the individual compound concentrations (14). For xenograft experiments, NBSGW mice were intravenously transplanted with MV4-11 AML cells (Fig. 4A). Two weeks after transplantation, engraftment of AML cells was confirmed (evaluated by gating on hCD45+ population by flow cytometry in peripheral blood). Treatment then began with vehicle, 50 mg/kg IMT, 100 mg/kg venetoclax, or a combination of IMT (50 mg/kg) and venetoclax (100 mg/kg) (Fig. 4A), dosed as such based on previously published protocols (14, 18). The efficacy of treatments was determined by measuring the proportion of transplanted AML cells in peripheral blood and by survival analysis (Fig. 4B). In all treatment groups, the hCD45+ population dropped immediately after treatment, and vehicle- and venetoclax-treated mice then exhibited a rapid rebound. In contrast, in IMT-treated mice the increase in leukemic cell proportion was delayed until eight weeks post-transplantation, and in the combination-treated group for to up to 20 weeks (Fig. 4B). These observations correlate with results from survival analysis (Fig. 4C). IMT treatment of MV4-11 transplanted mice significantly delayed the development of leukemia, inducing a 37% increase in median overall survival (57.5 → 79 days). Development of leukemia was further delayed by combination treatment to 131% (57.5 → 133 days), and the group contained two mice that did not show any disease symptoms after 200 days (post-transplantation) when they had to be euthanized according to ethical permissions. Moreover, leukemic cells were not detectable in the BM of surviving mice at the time of termination. All other mice showed symptoms of leukemia and of these, all but one combination-treated mouse had an increased percentage of leukemic cells in BM (Fig. 4D). These results confirm that treatment with a combination of IMT and venetoclax significantly prolongs leukemia progression in mice. Moreover, the general condition of mice such as behavior, fur, mobility and weight, assessed before (week 2) and after treatment (week 5), were not altered in the IMT-, venetoclax-, or combination-treatment groups when compared to the vehicle-treated group (weights shown in Fig. S5B), suggesting that the treatments were well tolerated. As with single IMT therapy (14), treatment with combination therapy did not display any evidence of organ toxicity or drug-associated death in the mice (Fig. S5C, D).

**Figure 4.**
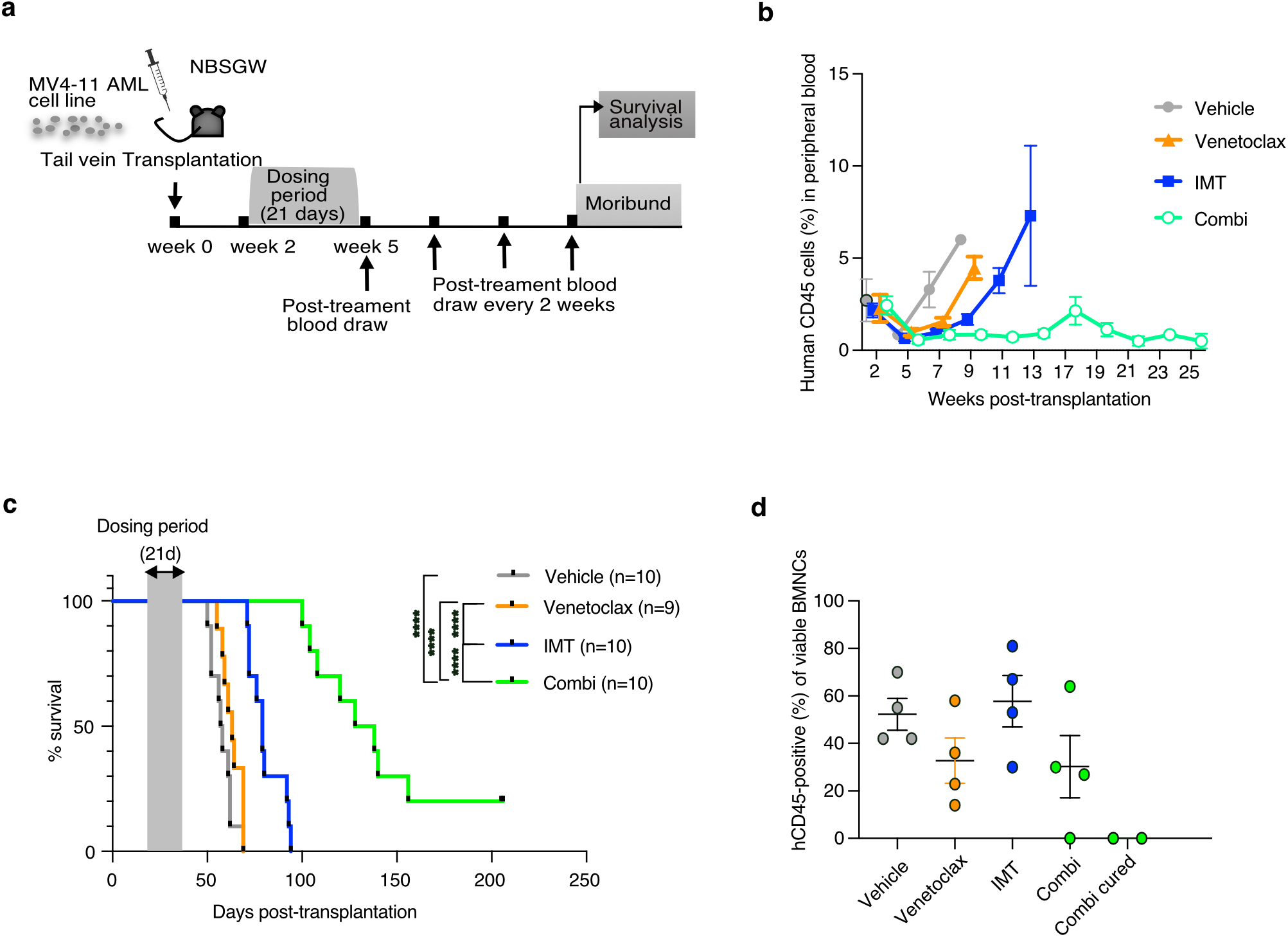
Combination of IMT and venetoclax significantly prolongs the survival of mice with leukemia. (**A**) Outline of CDX efficacy experiment: NBSGW mice were transplanted with MV4-11 human leukemia cells, then treated with either vehicle, IMT, venetoclax, or a combination of IMT and venetoclax (n=10 per treatment group). Peripheral blood was taken before treatment (week 2) and after treatment at regular intervals until disease progression was observed. (**B**) Analysis of proportion of human CD45+ cells in mice peripheral blood in all treatment groups at indicated time points, determined by flow cytometry. (**C**) Kaplan-Meier analysis of survival. Statistical significance was calculated using log-rank (Mantle-Cox) test. ****p < 0.0001, Error bars represent ± SEM. (**D**) Analysis of proportion of human CD45+ cells in the bone marrow of all treatment groups (four mice per group compared to two surviving mice) at time of euthanization.

### IMT and venetoclax synergistically decrease the tumor burden and increase the life span in an AML-PDX mouse model resistant to venetoclax

The CDX mouse model demonstrated IMT could sensitize AML cell lines to BCL-2 inhibitors *in vivo*. We next analyzed the effects of combination treatment on a more refined *in vivo* model by generating xenografts using AML patient-derived cells (AML PDX). We first tested an AML-PDX mouse model with resistance to venetoclax (AML sample harboring TP53 mutation). Two weeks post-transplantation with AML PDX cells, NSGS mice were orally treated with IMT, venetoclax, or a combination of the two for 21 days (Fig. 5A). The efficacy of the treatments was determined based on the proportion of human AML cells (determined by hCD45+ population). Six out of 10 mice in each group were euthanized immediately after the last compound administration for end-of-treatment analysis. We found that the combination of IMT and venetoclax resulted in significant inhibition of tumor burden in blood, BM, and spleen of treated mice (Fig. 5B). By examining histology sections of BM and spleen, we found that tissues from combination-treated animals contained less engraftment of leukemic blasts than single- and vehicle-treated mice (Fig. 5C). Accordingly, the frequencies of hCD45+ cells in the BM and spleens of the combination-treated group were largely reduced in comparison to other groups (Fig. 5D), and engrafted tumor cells in combination-treated PDX mice showed a lower proliferating percentage in comparison to single or vehicle treated groups (shown by KI67+/hCD45+ immunohistochemistry staining; Fig. 5E). This demonstrates a synergy between IMT and venetoclax in reducing tumor burden.

**Figure 5.**
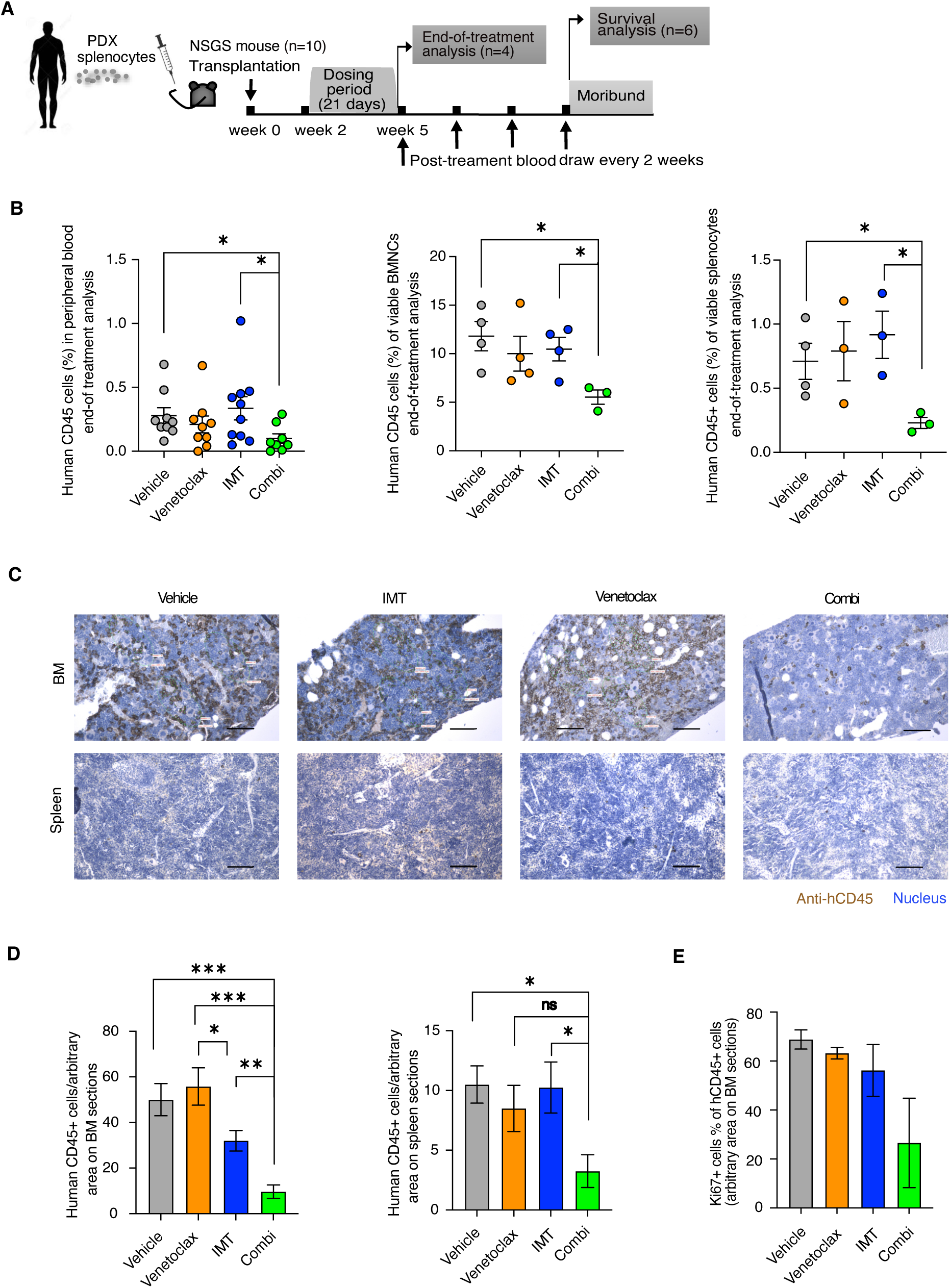
Synergistic effect of (IMT and venetoclax) combination treatment on tumor burden and survival of AML-PDX transplanted mice. (**A**) Experimental outline: NSGS mice were transplanted with primary AML splenocytes from PDX mice and were then treated with either vehicle, IMT, venetoclax, or a combination of IMT and venetoclax (n=10 per treatment group). After the final dosing, three or four mice were euthanized (end-of-treatment group) and the rest (n=6) were maintained for survival analysis. (**B**) Percentage of viable human CD45-positive (hCD45+) cells in peripheral blood, BM, and spleen of treated mice determined by flow cytometry. (**C**) Immunohistochemistry analysis of mouse BM and spleen sections stained with anti-human CD45 (brown) and counterstained with DAB (blue). (**D**) Quantitative analysis of hCD45+ cells on immunohistochemistry sections of BM and spleen sections of treated mice. (**E**) Proliferating tumor cells (CD45) are shown by quantification of Ki67 positive cells in immunohistochemistry-stained BM sections at end-of-treatment (results from at least three technical replicates and two biological replicates in each treatment group).

To further analyze the cellular effects of the IMT-venetoclax combination treatment, we analyzed transcriptomes of purified viable AML blasts isolated from PDX models at end of treatment (Fig. 6A). We confirmed that combination treatment had a more pronounced effect on gene transcription as compared to single treatments. In agreement with its more pronounced effect on tumor burden, combination treatment suppressed expression of several genes involved in leukemogenesis, including MX1, KCTD12, ITGA and RGL1 (Table S2) (19). Functional annotation and enrichment analysis were performed using the DAVID bioinformatics resources (Database for Annotation, Visualization, and Integrated Discovery) to identify significantly enriched gene ontology (GO) terms and pathways associated with the gene set. The gene list in Table S2 points to a potentially integrated function of the gene set in 1-Innate immune involvement, 2-Mitochondrial Function and Immune Signaling and 3-Signaling and Trafficking. The presence of annotations related to innate immune response, extracellular region, and transmembrane receptors suggests that the genes may be involved in the initial immune defense, recognizing pathogens, and activating immune cells. The indication of mitochondrial function and immune signaling approves that mitochondria are known not only for energy production but also for their emerging role in immune signaling, such as by producing reactive oxygen species (ROS) in response to infection. For signaling and trafficking, the presence of terms like “GTP binding,” “endosome,” and “calcium” highlights intracellular signaling, possibly in relation to immune activation or cellular responses The presence of terms like “GTP binding,” “endosome,” and “calcium” highlights intracellular signaling, possibly in relation to immune activation or cellular responses. We also analyzed survival by monitoring the mice for disease progression and morbidity. Peripheral blood engraftments were analyzed at regular intervals until the appearance of disease symptoms (Fig. 6B and Fig. S6A). The leukemia burden immediately prior to euthanization demonstrated that the cytoreductive effect of the combination treatment persists (Fig. 6B). However, blood count (determined by RBCs and HGB levels) did not vary significantly between experimental groups, nor did the magnitude of splenomegaly or liver weight vary at the time of euthanization (Fig. S6B–D). Interestingly, mice treated with a combination of IMT and venetoclax succumbed to death significantly later than other treatment groups, with a median survival of 112 days vs 96, 99, and 102 days for vehicle-, IMT-, and venetoclax-treated groups, respectively (Fig. 6**C**). Combination treatment thus increases the life span of venetoclax-resistant AML mouse models, demonstrating that IMT can sensitize AML cells towards other therapies.

**Figure 6.**
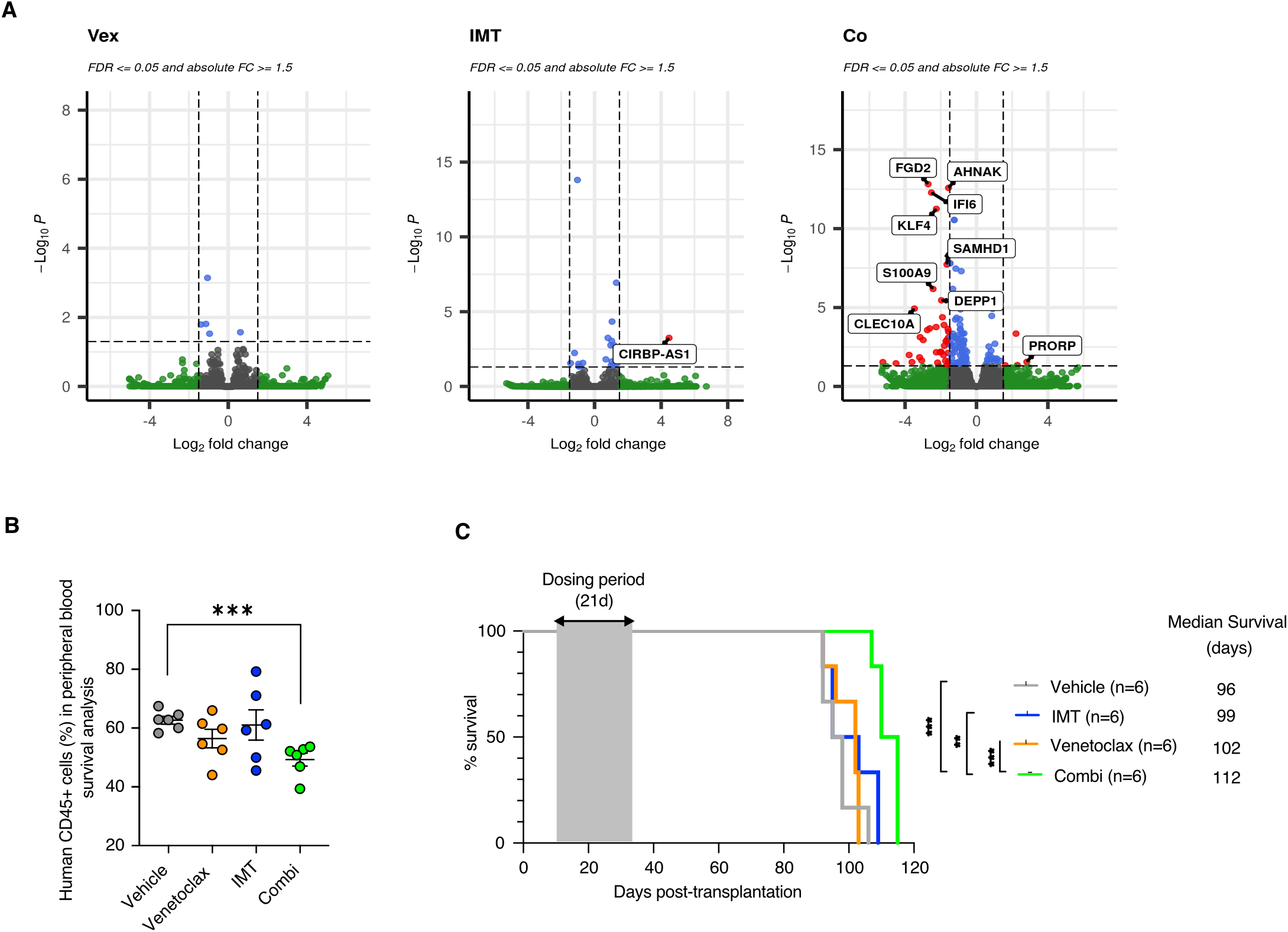
Synergistic effect of (IMT and venetoclax) combination treatment on gene transcription and survival of AML-PDX transplanted mice. The experimental outline was shown in figure 5A, but n=10 per treatment group. After the final dosing, three or four mice were euthanized (end-of-treatment group) and the rest (n=6) were maintained for survival analysis. (**A**) Gene expression profiling by RNA-seq of purified AML blasts isolated from PDX models treated with venetoclax, IMT, or a combination thereof. Indicated expression changes are relative to a vehicle treated group. Significantly enriched genes are indicated in red (q < 0.05). (**B**) Peripheral blood engraftment analysis of survival group. (**C**) Survival analysis of treated mice shown by Kaplan-Meier analysis (n=6/treatment group). *P < 0.05, **P < 0.01, and ***p < 0.001.

### IMT does not affect tumor burden or life span in an AML-PDX mouse model sensitive to venetoclax

For comparison, we also monitored effects of combination treatment on a PDX model based on AML cells sensitive to venetoclax treatment (AML cells harboring FLT3-ITD, NMP1 and DNMT3A). As expected, we observed that venetoclax treatment significantly reduced the tumor burden in the blood and BM of mice. Combination treatment with IMT did not add to this effect (Fig. 7A). By examining histology sections, we found that both BM and spleen tissues from combination- and venetoclax-treated animals contained reduced proportion of leukemic blasts compared to IMT- and vehicle-treated mice (Fig. 7B). White blood counts (WBCs), red blood counts (RBCs), and hemoglobin (HGB) levels were not significantly different across experimental groups (Fig. S7A). The total body and liver weight of PDX mice were very similar in all groups (Fig. S7B, D), but both venetoclax and combination treatments reduced splenomegaly (Fig. S7C).

**Figure 7.**
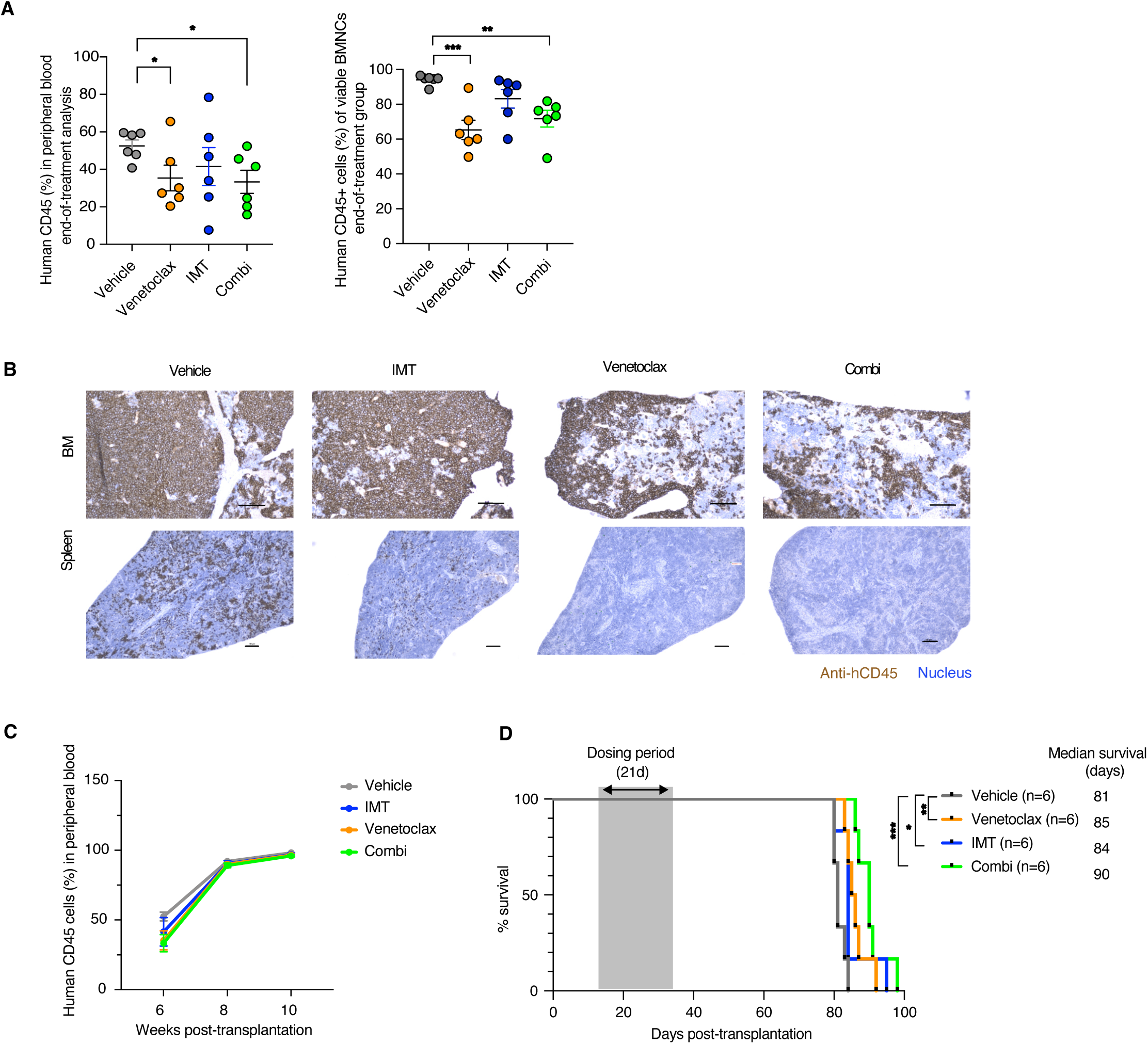
Leukemia progression induced by primary AML cells is inhibited by venetoclax treatment. As in Fig. 5A, efficacy experiment outline: NSGS mice were transplanted with primary AML splenocytes from PDX mice and were then treated with either vehicle, IMT, venetoclax, or a combination of IMT and venetoclax (n=12 per treatment group). 8–12 h after the last dosing, half of the mice (n=6/treatment group) were euthanized (end-of-treatment group) and the other half were maintained for survival analysis. (**A**) Percentage of viable human CD45-positive (hCD45+) cells in peripheral blood and bone marrow of treated mice (n=6 for each treatment group), determined by flow cytometry. (**B**) Immunohistochemistry analysis of hCD45+ cells on bone marrow and spleen tissue sections of treated mice (images are representative from at least two mice per group). Scale bars represent 100 μM. (**C**) Peripheral blood engraftment analysis of survival group. (**D**) Mice treated with single or a combination of compounds showed prolonged survival in comparison to the vehicle-treated group. Kaplan-Meier analysis shows the relative survival curve. *P < 0.05, **P < 0.01, and ***p < 0.001.

The remaining mice were used for survival analysis, which involved regular blood count and chimerism analysis until the appearance of disease symptoms (Fig. 7C). Both venetoclax and IMT caused a modest increase in life span, but combination treatment did not offer an additional effect (Fig. 7D).

## Discussion

In recent years, the genetic complexity and heterogeneity of leukemia cell populations has become apparent (20, 21) and developing effective strategies to target human leukemia initiating cells has thus proven to be a challenging task. Currently, there is considerable interest in therapies, such as venetoclax, that can restore normal apoptosis of cancer cells by directly interacting with the BCL-2 family of proteins. These are effective for some leukemias, but many patients eventually become refractory to treatment and ultimately succumb to their disease (22, 23). It has become increasingly important to determine which biological properties are relatively consistent in leukemic cell populations irrespective of genetic makeup (tumor population). Recent observations indicate that several cancer cell types, including AML, are reliant on OXPHOS for metabolism (24), making inhibition of mitochondrial function a promising therapeutic strategy for AML (12, 25–28). Therapy-resistant leukemic cells are more reliant on oxidative respiration than glycolysis for energy generation, suggesting that the OXPHOS dependency and lack of normal glycolytic activity in AML cells may be characteristic of cancers (29). Understanding resistance mechanisms will allow for the design of strategies to counteract drug resistance or impede resistance from developing. In AMLs with aberrant expression of BCL-2, venetoclax can reduce OXPHOS (9) and trigger cell death. However, venetoclax resistance can occur in patients treated with the BCL-2 inhibitor and in some subtypes of AML patients with poor prognoses (30).

Previous work has indicated that AML cells display increased mitochondrial biogenesis and inhibition of mitochondrial translation can be used as a strategy for human AML (18, 27). In our work, we target another process involved in mitochondrial gene expression, i.e., transcription. IMT treatment targets POLRMT, which has two main functions in mitochondria: it transcribes mtDNA encoded genes and provides the primers required to initiate mtDNA replication. Inhibition of POLRMT can thus deliver a double punch, both reducing mtDNA copy number and inhibit OXPHOS biogenesis. As demonstrated here inhibition of POLRMT can overcome venetoclax therapy resistance in AML. These observations are in line with recent reports of the importance of mitochondrial function in leukemia cell survival (31, 32). By developing a clinically relevant and well-tolerated approach to deliver IMTs in immunodeficient mice transplanted with AML cells, we were able to reduce the tumor burden in all (cell- and patient-derived) xenografted mice treated with a combination of IMT and venetoclax. This demonstrates that reducing POLRMT activity results in successful targeting of leukemic cells. Our *in vitro* investigations confirmed that the combination of IMT and venetoclax is necessary to significantly decrease ATP levels and consequently induce apoptosis. In addition, efficacy studies of AML PDX models showed that single compounds are not sufficient to eradicate AML cells, but that the combination regimen reduced AML tumor burden and prolonged survival in all treated mice. These data indicate the advantage of both downregulating POLRMT activity in combination with increasing apoptosis and highlights the importance of rational combination therapies (33, 34). IMT offers potential advantages over other OXPHOS-targeting therapies due to its lower toxicity, as observed in preclinical models. While other strategies, such as mitochondrial translation inhibition or electron transport chain inhibitors, effectively target OXPHOS in AML, they may come with higher toxicity risks. IMT’s selective targeting of mitochondrial transcription appears to minimize adverse effects, making it a promising candidate for use in combination regimens with BCL-2 inhibitors like venetoclax. This approach could improve therapeutic outcomes while maintaining tolerability, addressing a significant challenge in AML treatment.

In our study, we observed that the combination of IMT and venetoclax did not show synergistic effects in TP53 mutant AML cells, particularly in the THP-1 cell line. Although statistical analyses indicated potential interaction, the biological impact was minimal in TP53 mutant cells, contrasting with the robust synergy observed in TP53 wild-type cells like MV4-11 and MOLM-13. This suggests that TP53 mutation status may influence the efficacy of this combination therapy, highlighting a potential limitation in targeting mitochondrial transcription to potentiate BCL-2 inhibition in TP53 mutant AML context. These findings underscore the need for broad mutational profiling when designing combination therapies for AML.

In summary, we have introduced a new therapeutic strategy to target therapy-resistant leukemia with OXPHOS-dependent properties. Despite several limitations with the design of pre-clinical cancer models and the intensity and duration of treatments, we demonstrate here that the combined inhibition of BCL-2 and mitochondrial transcription can target therapy-resistant AMLs. This combination treatment succeeds as IMT primes cell vulnerability to apoptosis induction. Consequently, our findings may be of value for the future development of better clinical strategies for the treatment of AML.

## Acknowledgments

The authors would like to thank the staff at the Department of Experimental Biomedicine at the University of Gothenburg, namely, Jenny Nyman, Claudia Engwall, Tova Johansson and Hanna Brissman for tremendous assistance with mouse experiments. They would also like to thank Dr Rebekah Hailes for editing the manuscript and to acknowledge Lenka Palova-Jelinkova, Klara Danova, and Pavla Jestrabova from SOTIO Biotech (Prague, Czech Republic) for valuable input on *in vitro* and *in vivo* studies. The authors acknowledge support from the National Genomics Infrastructure in Stockholm funded by Science for Life Laboratory, the Knut and Alice Wallenberg Foundation and the Swedish Research Council, and SNIC/Uppsala Multidisciplinary Center for Advanced Computational Science for assistance with massively parallel sequencing and access to the UPPMAX computational infrastructure. This research is supported by Swedish Research Council (2017-01257:CMG), Swedish Cancer Foundation (20 1291 PjF:CMG, and 20 0925 PjF:LP), Knut and Alice Wallenberg Foundation (2017.0080: CMG), The Swedish state under the agreement between the Swedish government and the county councils, the ALF agreement (ALFGBG-966275:CMG and ALFGBG-965206: LP), and Assar Gabrielsson Foundation for cancer research grant (LSA)

## Materials and Methods

### Cell lines and culture conditions

All cell lines (MV4-11, MOLM13, THP-1, KG-1, OCI-AML3, and OCI-AML2) were purchased from DSMZ (Braunschweig, Germany) and cultured in RPMI-1640 (Sigma Aldrich) supplemented with 10–20% heat inactivated fetal bovine serum (Gibco) and 1% penicillin/streptomycin (Invitrogen). Cells were maintained in a humidified incubator at 37 °C and 5% CO_2_. RPMI-1640 without glucose was used for *in vitro* treatment experiments and supplemented by physiological concentrations (5 mM) of glucose (D-Glucose, Sigma) or galactose (D-galactose, Sigma).

### Generation of *in vivo* AML mouse model

Mice were purchased from Jackson laboratories (USA) and maintained in a pathogen-free facility at the University of Gothenburg. All animal experiments were conducted in accordance with guidelines approved by the research animal ethics committee at the University of Gothenburg. For AML cell line xenograft (CDX) experiments, 6–8-week-old, male NBSGW mice (NOD.Cg-Kit^W-41J^ Tyr^+^ Prkdc^scid^ Il2rg^tm1Wjl^/ThomJ, Jackson laboratories) were used. Mice were transplanted intravenously with ∼1 × 10^6^ MV4-11 cells. For patient-derived xenograft (PDX) experiments, 6–8-week-old, male NSGS mice (NOD.Cg-Prkdc^scid^ Il2rg^tm1Wjl^ Tg (CMV-IL3,CSF2,KITLG)1Eav/MloySzJ, Jackson laboratories) were sub-lethally irradiated with 1 Gy radiation. AML PDX splenocytes were purchased from Jackson laboratories (id: J000106134*, male, 58 years old, AML, M4, positive for FLT3-ITD, DNMT3A, and NPM1mutaions; and id: 000106566*, male, 48 years old, AML no classification specified, positive for TP53 mutation). To obtain adequate number of PDX cells for the scale of the efficacy experiments, the splenocytes were initially expanded in vivo. Mice were transplanted intravenously with ∼1 × 10^6^ splenocytes from AML PDX mice containing more than 80% patient AML cells.

### Human leukemia cell frequency assessment by flow cytometry

The percentage of human leukemic cells (MV4-11 cell lines and AML PDX cells) in mouse peripheral blood, BM, and spleen was determined by flow cytometry. Mice were bled at indicated time points after transplantation. Prior to staining, peripheral red blood cells were lysed using ammonium chloride for 8–10 min at room temperature. Cells were stained with antibodies, washed, and submitted for flow-cytometric analysis using a FACS Aria (BD Biosciences, USA) or CytoFlex instrument (Beckman Coulter, California, USA). At least 10,000 events were acquired, and the percentage was calculated relative to gated viable mononuclear cells (MNCs). To discriminate between human and mouse cells, cells were stained with antibodies against human CD45 (clone HI30, BD Biosciences) and mouse CD45 (clone 2D1, BD Biosciences). Engraftment was deemed positive if human CD45+ cells represented >0.1% of the viable MNCs. 7AAD (BD Biosciences) served as viability marker.

### Preparation of compounds for *in vivo* drug administration

IMT was optimized based on the structure activity and property relationship of the coumarin mtRNAPol inhibitor class by LDC (Dortmund, Germany). Venetoclax was obtained from MedChemExpress (Hölzel Diagnostika Handels GmbH, Germany). Following the development of a suitable vehicle for the formulation of IMT and venetoclax, as well as the combination of both compounds, the test articles were formulated using 0.5% aqueous methylcellulose solution (Sigma-Aldrich, Germany). After randomization, and two weeks post-transplantation or with confirmation of engraftment, mice were treated with vehicle (0.5% methylcellulose), 50 mg/kg IMT, 100 mg/kg venetoclax, or a combination (100 mg/kg venetoclax + 50 mg/kg IMT) by oral administration daily for 21 days.

### Cell proliferation assay

Cells were seeded out ca 20 h prior to treatment. For dose response experiments, cells were treated with a wide range of concentrations of single compounds. Cell proliferation assays were performed every 24 h from 72–168 h, where cell viability/ATP production was determined using CellTiter-Glo (Promega).

### Apoptosis assay and Mitochondrial potential assay

Apoptosis was evaluated by Annexin V and PI staining and mitochondrial membrane potential was evaluated by lipophilic cationic dye JC-1. Cells were treated with IC50 concentrations of IMT, venetoclax, or combination of both, analyzed using an advanced image cytometer (Nucleocounter NucleoCounter’s accompanying software NucleoView).

### High-resolution respirometry

AML cell lines were treated with DMSO, IMT, venetoclax, or combination of IMT and venetoclax for 3–4 days. The mitochondrial respiratory capacity (oxygen consumption rate) was measured using high resolution respirometry (O2K, Oroboros, Innsbruck, Austria). A substrate-uncoupler-inhibitor titration (SUIT) protocol was used to measure specific features of mitochondrial respiration. Respiratory capacities were tested in a sequence of coupling states: basal respiration; LEAK respiration using 5 mM oligomycin (Sigma); and non-coupled electron transfer state (ET state) where the optimum carbonyl cyanide 3-chlorophenylhydrazone (CCCP, Sigma) concentration for maximum flux was determined by titration (max. 3 μM CCCP). Mitochondrial respiration was inhibited using 2.5 μM antimycin (Sigma). Where stated, excess glucose (D-Glucose, Merck) was added at a final concentration of 20 mM. Instrumental and oxygen background fluxes were calibrated as a function of oxygen concentration and subtracted from the total volume-specific oxygen flux. Calibration and data analysis was performed using Datlab v.6.1 software (Oroboros Instruments).

### Pharmacokinetics analysis

IMT and venetoclax were extracted from plasma and tissues by protein precipitation using acetonitrile. Tissue samples were homogenized using double the (sample) volume of PBS, and bone marrow samples were suspended in 40 µL of PBS prior to extraction with acetonitrile. Samples were analyzed by liquid chromatography tandem-mass spectrometry using a Prominence UFLC system (Shimadzu) coupled to a Qtrap 5500 instrument (ABSciex). Test articles were separated on a C18 column using a gradient elution with an acetonitrile/water mixture containing 0.1% formic acid as the mobile phase. Chromatographic conditions and mass spectrometry parameters were optimized for each test article prior to sample analysis. Concentrations of each test article were calculated by means of a standard curve.

### mRNA extraction and transcriptome sequencing of AML PDX cells

For RNAseq experiment BM samples were FACS sorted using BD FACS Aria fusion (BD Biosciences). Human AML cells were isolated using human CD45 antibody. 7-AAD (BD Biosciences) was used for dead cell exclusion. RNA was extracted using RNeasy Micro kit (Qiagen). RNAseq library preparation and RNA-sequencing was done using the SMARTer Total Stranded RNA pico kit. RNA library was prepared at National Genomics Infrastructure (NGI) to be sequenced on the NovaSeq S6000 on 0.5 lanes of one S4 flowcell.

### Bioinformatics analysis

Analysis was performed in R (www.r-project.org, accessed on 10 May 2022) using DESeq2, clusterProfiler and ReactomePA packages. Prefiltering to keep genes that have at least 10 reads total was performed before differential expression analysis was done with DESeq2. Visualization of differentially expressed genes was done with EnhancedVolcano to create volcano plots and gridExtra for creating tables. The *p*-value cutoff was set to 0.1 to allow for more genes to be found in EP (End-of-treatment) group comparisons and the Res/Sen (Resistant/Sensitive) group comparisons. Upregulated and downregulated genes were plotted and colored according to Fold Change (FC) greater than 1.5 or -1.5 and the *p*-value cutoff 0.1.

Functional annotation and enrichment analysis were performed using the DAVID bioinformatics resources (Database for Annotation, Visualization, and Integrated Discovery) to identify significantly enriched gene ontology (GO) terms and pathways associated with the gene set (35).

### Statistical Analysis

Unless otherwise noted, data were summarized using mean ± standard error of the mean (± SEM). Per-group sample sizes presented in figures and results are reported from ≥3 separate experiments, unless stated otherwise. Survival distributions were estimated using the Kaplan– Meier method, and group comparisons were conducted using the log-rank test. Data were analyzed using GraphPad Prism 9.0. Combination analysis was summarized using the Bliss-independent model (16). The formula Y_ab,P_ = Y_a_ +Y_b_ − Y_a_Y_b_ was used where drug A at dose a inhibits Y_a_ percent of cell growth, and drug B at dose b inhibits Y_b_ percent of cell growth. The predicted inhibitory effect of combined percentage was estimated by Y_ab,P_. The observed combined percentage inhibition Y_ab,O_ was then compared with Y_ab,P_, and if Y_ab,O_ > Y_ab,P_, the combination treatment was thought to be more efficacious (synergy). If Y_ab,O_ = Y_ab,P_ the combination treatment was thought to be additive.

## Author contributions

Conceptualization (LSA, RD, TB, AU, PN, BMK)

Methodology (TB, AU, PN, BMK)

Investigation (LSA, JA, CN)

Supervision (BMK, LP, CG)

Writing—original draft (LSA)

Writing—review & editing (LSA, JA, AU, RD, TB, PN, BMK, LP, CMG)

## Competing interests

CMG is a scientific co-founder of Pretzel Therapeutics Inc. Together with RD, TB, AU, PN, BMK who are employees of LDC, CMG is also co-inventor of the patent application WO 2019/057821. The Medicinal chemistry and pharmacology part of this work was financed by the Max-Planck Gesellschaft e.V. under the framework agreement between Max-Planck and LDC.

## Data and materials availability

All data needed to evaluate the conclusions in the paper are present in the paper and/or the Supplementary Materials. RNA-seq datasets have been deposited in the GSE repository (GSE213976). Additional data related to this paper may be requested from the authors.

## Notes

### Summary of Updates

This version of the manuscript has been revised based on reviewer feedback from our previous submission to eLife. Key updates include: Pathway Analysis: We have incorporated gene ontology (GO) analysis of transcriptomic data, highlighting pathways involved in mitochondrial function, immune signaling, and intracellular signaling, providing mechanistic insight into the drug combination. Synergy Analysis: A Bliss synergy matrix has been added to support the synergistic interaction between IMT and venetoclax, demonstrating enhanced inhibition of AML cell viability. Statistical Comparisons: We have included statistical analyses comparing venetoclax alone versus combination treatment (Fig. 5C) and clarified statistical methods for OXPHOS measurements. PDX Model Characterization: Additional details on the classification and genetic background of patient-derived xenograft (PDX) models have been included for better context. Bone Marrow Sanctuary Hypothesis: We now discuss the potential for leukemic cell persistence in the bone marrow, despite the absence of leukemia in peripheral blood, providing a rationale for disease relapse. Limitations & Future Directions: While our study focuses on mitochondrial inhibition, we acknowledge the importance of glycolysis measurements, liver and kidney toxicity assessments, and alternative AML therapy combinations, which we suggest for future research. These revisions enhance the manuscript's clarity and provide a more comprehensive interpretation of our findings.

