## Supplementary Materials for "Reduced mitochondrial transcription sensitizes acute myeloid leukemia cells to BCL-2 inhibition"

†Lead Contact

**Supplementary Materials**

**Materials and Methods**

**General Chemistry for compound synthesis**

Anhydrous solvents and reagents were purchased from various fine chemical suppliers and were used without further purification. ^1^H NMR spectra were taken on a Oxford Varian 400/54 (400 MHz) spectrometer or a Bruker Avance II (300 MHz) with residual protonated solvent (CHCl_3_ δ 7.26; DMSO δ 2.49) as reference. Data was reported as follows: chemical shift, multiplicity (bs, broad singlet; s, singlet; d, doublet; t, triplet; q, quartet; and m, multiplet), coupling constant, and integration. ^13^C NMR spectra were taken on a Brucker Biospin with power gated H1 decoupling (150 MHz) spectrometer and showed an unusually complicated splitting pattern, due to the presence of diasteromers, Carbon-Fluorine Coupling and rotational effects. For this reason, the peaks assigned to the same carbon are shown between brackets. Final compounds were judged to be ≥95% pure by HPLC analysis using a Waters Acquity UPLC (Sample Manager, Binary Solvent Manager, Column Heater/Cooler, PDA eλ Detector; SQ Detector), Column Acquity UPLC BEH C18 1.7 µm, 2.1 x 50 mm, eluted with H_2_O with 0.05% formic acid (Solvent A) and MeCN with 0.05% formic acid (Solvent B) at 0.5 mL/min. Gradient: elution from 5% to 100% B over 3.5 min with an initial hold of 0.5 min and a final hold at 100% B of 0.5 min. Total run time: 5 min or a HPLC/MS Waters (2767 Sample Manager, 515 HPLC Pump, 2525 Binary Gradient Module, 2996 Photodiode array Detector, Micromass ZQ Detector), Column Xterra® MS C18 5μm 100 x 4.6mm, eluted with H_2_O with 0.1% formic acid (Solvent A) and MeCN (Solvent B) at 2 mL/min. Gradient: elution from 5% to 100% B over 7 min and a final hold at 100% B of 1.5 min. Total run time: 8.5 min. Reported yields are not optimized, with emphasis on purity of products rather than quantity.

**Condensation reaction to obtain intermediate 1**

Ethyl 3-(2-chloro-4-fluorophenyl)-3-oxopropanoate (1)

60% NaH in mineral oil (4.86 g, 121.7 mmol) was washed with pentane and dried under a flow of nitrogen. Dry toluene (350 mL) was added and the suspension was cooled to 0°C. Diethyl carbonate (27.4 g, 231.8 mmol) was added dropwise over a period of 25 min, followed by 2-chloro-4-fluoroacetophenone (10 g, 57.9 mmol) over a period of 20 min. The cooling bath was removed, and the reaction heated to 50°C and stirred for 18 h. The reaction mixture was allowed to cool to r.t. and was poured into ice cold water (500 mL). The aqueous layer was acidified to pH 2 with 10% aq. HCl and extracted with Et_2_O. The combined organic phases were dried over MgSO_4_, filtered and evaporated *in vacuo* to yield the title compound 1b (6.0 g, 42%) as a mixture of tautomers, which was used in the following step without further purification. Tautomer 1, ethyl 3-(2-chloro-4-fluorophenyl)-3-oxopropanoate: ^1^H NMR (300MHz, Chloroform-d) δ 7.63 (dd, J = 8.8, 6.0 Hz, 1H), 7.14-6.93 (m, 2H), 4.12 (q, J = 7.1 Hz, 2H), 3.96 (s, 2H), 1.18 (t, J = 7.1 Hz, 3H); LCMS tR = 5.18 min, MS (ESI+) m/z 245.19, 247.18 (M+H)^+^, 43%. Tautomer 2, (Z)-ethyl 3-(2-chloro-4-fluorophenyl)-3-hydroxyacrylate: ^1^H NMR (300MHz, Chloroform-d) δ 12.43 (s, 1H), 7.52 (dd, J = 8.7, 6.1 Hz, 1H), 7.14-6.93 (m, 2H), 5.48 (s, 1H), 4.21 (q, J = 7.1 Hz, 2H), 1.27 (t, J = 7.1 Hz, 3H); LCMS tR = 6.38 min, MS (ESI+) m/z 245.19, 247.18 (M+H)^+^, 34%.

**Pechmann reaction to obtain intermediate 2**

4-(2-Chloro-4-fluorophenyl)-7-hydroxy-2H-chromen-2-one (2)

Intermediate 1 (28.2 g, 115 mmol) was dissolved in methansulfonic acid (850 mL) and resorcinol (12 g, 109 mmol) was added. The mixture was stirred at 45°C for 2 h, upon which the reaction was allowed to cool to r.t. EtOH was added dropwise, followed by water. The suspension was extracted with EtOAc and washed with brine. The combined organic layers were dried over MgSO_4_, filtered, and evaporated *in vacuo*. The crude product was purified by flash chromatography on silica gel using a gradient of EtOAc in cHex to yield the title compound 2 (25.4 g, 80%) as a pink solid. ^1^H NMR (300 MHz, DMSO-d6) δ 10.70 (bs, 1H), 7.69 (dd, J = 8.9, 2.5 Hz, 1H), 7.55 (dd, J = 8.6, 6.1 Hz, 1H), 7.42 (td, J = 8.5, 2.6 Hz, 1H), 6.91 – 6.77 (m, 2H), 6.74 (dd, J = 8.7, 2.3 Hz, 1H), 6.21 (s, 1H); LCMS tR = 5.28 min, MS (ESI+) m/z: 291.10, 293.16 (M+H)^+^, 95%.

**Mitsunobu reaction to obtain intermediate 3**

(R)-Ethyl 2-((4-(2-chloro-4-fluorophenyl)-2-oxo-2H-chromen-7-yl)oxy)propanoate (3)

Intermediate 2 (12 g, 41.28 mmol) and PPh_3_ (11.91 mg, 45.41 mmol) were dissolved in THF (7 mL), and (-)-Ethyl (S)-2-hydroxypropionate (7.07 ml, 61.92 mmol) was added. The reaction was cooled to 0°C and DIAD (8.94 ml, 45.41 mmol) was added dropwise. The reaction was then stirred at r.t. for 4 h. The mixture was diluted with EtOAc and washed with a sat. NaHCO_3_ solution, a sat. NH_4_Cl solution, and water. The organic layer was dried over MgSO_4_, filtered, and evaporated *in vacuo*. The crude product was purified by flash chromatography on silica gel using a gradient of EtOAc in cHex to yield the title compound 3 (13.53 g, 83%) as a colorless glue. ^1^H NMR (400MHz, DMSO-d6) δ 7.70 (ddd, J = 8.9, 2.5, 1.5 Hz, 1H), 7.57 (ddd, J = 8.6, 6.1, 1.5 Hz, 1H), 7.43 (tdd, J = 8.5, 2.6, 1.3 Hz, 1H), 7.02 (dd, J = 5.9, 2.4 Hz, 1H), 6.95 (dd, J = 8.9, 2.4 Hz, 1H), 6.88 (ddd, J = 8.9, 3.6, 2.5 Hz, 1H), 6.33 (s, 1H), 5.18 (dd, J = 6.8, 1.1 Hz, 1H), 4.16 (qd, J = 7.1, 1.4 Hz, 2H), 1.54 (dd, J = 6.8, 0.5 Hz, 3H), 1.19 (t, J = 7.1 Hz, 3H); LCMS tR = 3.23min, MS (ESI+) m/z: 391.29, 393.32 (M+H)^+^, 100%.

**Hydrolysis reaction to obtain intermediate 4**

(R)-2-((4-(2-chloro-4-fluorophenyl)-2-oxo-2H-chromen-7-yl)oxy)propanoic acid (4)

Intermediate 3 (13.53 g, 34.62 mmol) was dissolved in THF and 2 M NaOH aq. solution (250 mL) was added at 0°C. A few drops of MeOH were added until the mixture was homogeneous. The reaction was allowed to warm up to r.t. stirring for 30 min, neutralized with 2 M HCl, and stirred for a further 30 min. The mixture was extracted with EtOAc and the combined organic phases were washed with H_2_O, dried over MgSO_4_, filtered, and evaporated *in vacuo* to yield the title compound 4 (12.55 g, 100%) as a white solid. ^1^H NMR (400 MHz, DMSO-d6) δ 13.21 (s, 1H), 7.68 (ddd, J = 8.9, 2.5, 1.9 Hz, 1H), 7.54 (dd, J = 8.6, 6.1 Hz, 1H), 7.46 – 7.37 (m, 1H), 7.00 – 6.89 (m, 2H), 6.85 (ddd, J = 8.9, 2.5, 1.5 Hz, 1H), 5.04 (qd, J = 6.8, 2.0 Hz, 1H), 1.51 (d, J = 6.7 Hz, 3H); LCMS tR = 2.70min, MS (ESI+) m/z: 363.30, 365.25 (M+H)^+^, 100%.

**Amide coupling to obtain intermediates 5**

Ethyl 2-((R)-1-((R)-2-((4-(2-chloro-4-fluorophenyl)-2-oxo-2H-chromen-7-yl)oxy)propanoyl)piperidin-3-yl)acetate (5)

Intermediate 4 (10.52 g, 29.00 mmol) EDC.HCl (8.34 g, 43.50 mmol), and HOBt.xH2O (6.71 g, 43.80 mmol) were dissolved in DMF (300 mL) and (R)-Piperidin-3-yl-acetic acid ethyl ester hydrochloride (6.08g, 29.29 mmol) was added slowly. Et_3_N (6.06 ml, 43.50 mmol) was added, and the mixture was stirred at r.t. for 1 h. The reaction was diluted with EtOAc and washed with H_2_O, a sat. NaHCO_3_ solution, a sat. NH_4_Cl solution, and again with H_2_O. The organic phase was dried over MgSO4, filtered and evaporated in vacuo. The crude product was purified by flash chromatography on silica gel using a gradient of EtOAc in cHex to yield the title compound 5 (11.93g, 80%) as a colorless foam. ^1^H NMR (400 MHz, DMSO-d6) δ 7.69 (dd, 1H), 7.61–7.51 (m, 1H), 7.43 (dddd, 1H), 7.00–6.89 (m, 2H), 6.86–6.78 (m, 1H), 6.31 (s, 1H), 5.59–5.25 (m, 1H), 4.15 (t, 1H), 4.09–4.03 (m, 2H), 3.89 (dd, 1H), 3.15–2.89 (m, 1H), 2.66–2.52 (m, 1H), 2.34–2.12 (m, 2H), 1.78 (s, 2H), 1.64 (s, 1H), 1.52–1.37 (m, 3H), 1.25 (s, 2H), 1.19–1.13 (m, 3H); LCMS tR = 3.12min, MS (ESI+) m/z: 516.35, 518.38 (M+H)^+^, 100%.

**Hydrolysis reaction to obtain final compound LDC204857**

2-((R)-1-((R)-2-((4-(2-chloro-4-fluorophenyl)-2-oxo-2H-chromen-7-yl)oxy)propanoyl)piperidin-3-yl)acetic acid (LDC204857)

Intermediate 5 (11.53g, 22.35mmol) was dissolved in THF (335ml), and a 2 M NaOH aq. solution (67 mL) was added at 0°C. The reaction was stirred at r.t. for 1 h, acidified to pH 3 with 2M HCl, and stirred for further 30 min. The mixture was extracted with EtOAc and the combined organic phases were washed with water, dried over MgSO_4_, filtered and evaporated *in vacuo*. The crude was purified by reverse phase flash chromatography on C18 using a gradient of MeCN in H_2_O (with 0.5% formic acid) to yield the title compound LDC204857 (9.90 g, 91%) as a white solid. m.p.: 109.4°C; ^1^H NMR (400 MHz, DMSO-d6) δ 12.12 (br. s, 1H), 7.69–7.62 (m, 1H), 7.61–7.43 (m, 3H), 6.93–6.85 (m, 2H), 6.84–6.75 (m, 1H), 6.27 (s, 1H), 5.44-5.35 (m, 1H), 4.16 (dd, 1H), 3.88 (dd, 1H), 3.20–2.70 (m, 1H), 2.61-2.52 (m, 1H), 2.56-2.37 (m, 1H), 2.21– 2.06 (m, 2H), 1.94–1.57 (m, 2H), 1.44 (t, 2H), 1.36–1.19 (m, 3H); ^13^C NMR (101 MHz, DMSO-d6) δ [173.60, 173.49], [168.08, 167.96], 164.06, [161.57, 161.55], [161.17, 161.08], [160.22, 160.19], [155.25, 155.20], [152.62, 152.58], [132.65,132.88, 132.65, 132,55], [130.79, 130.76], [128.24, 127.98], [117.78, 117.53], [115.63, 115.42], [113.71, 113.67, 113.64, 113.59], 102.44, 71.56, 50.15, 47.11, 45.47, [38.19, 37.97], 33.86, 33.07, 30.43, 25.65, 24.88, [18.04, 17.81].; LCMS tR = 2.67 min, MS (ESI+) m/z: 488.29, 490.25 (M+H)^+^, 100%.

**Supplementary Figure S1.**

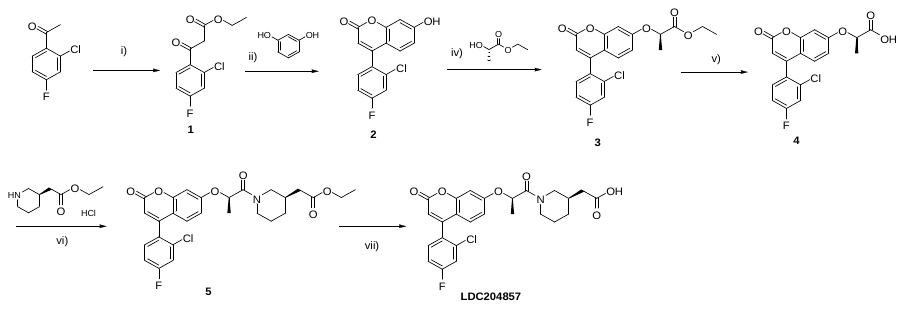

**Fig. S1.** Synthesis scheme of compound LDC204857. i) (EtCO)_2_O, 60% NaH in mineral oil, toluene, 50°C, 18h; ii) MsOH, 45°C, 2h; iii) Cs_2_CO_3_, DMF, r.t., 1.5h; iv) PPh_3_, DIAD, THF, 0°C to r.t., 4h; v) 2M NaOH, THF, 0°C to r.t., 30 min, b) 2M HCl, 30min; vi) EDC.HCl, HOBt.xH2O, Et_3_N, DMF, r.t., 1.5h; vii) a) 2M NaOH, THF, MeOH, 0°C to r.t., 1h, b) 2M HCl, 30min.

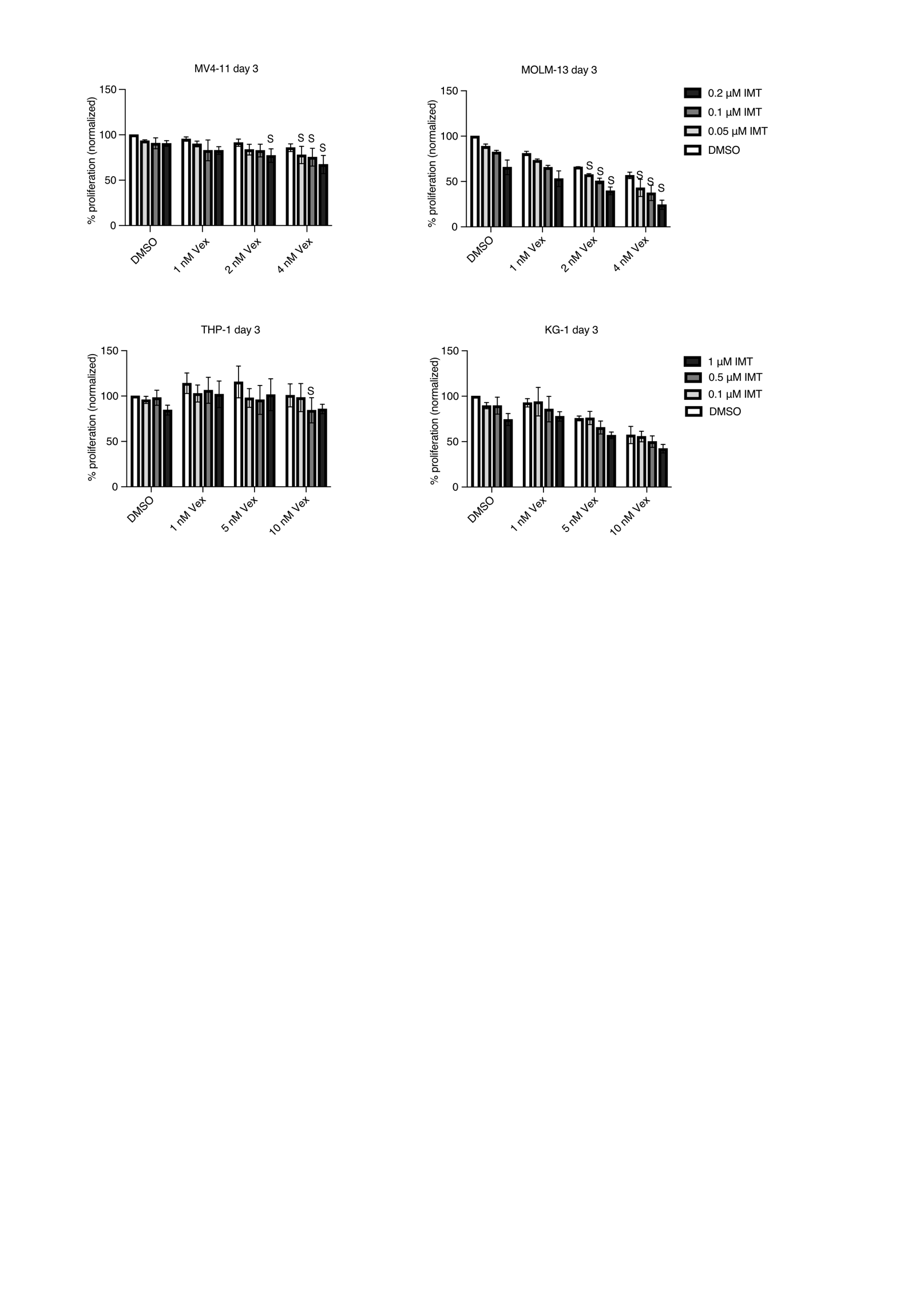

**Figure S2.** Synergism analysis of AML cell lines treated with a combination of IMT (0.05, 0.1, 0.2 μM for MV4-11 and MOLM-13, and 0.1, 0.5, 1 μM for THP-1 and KG-1) and venetoclax (Vex; 1, 2, 4 nM for MV4-11 and MOLM-13, and 1, 5, 10 nM for THP-1 and KG-1) for three days.values are shown as percentage proliferation normalized to DMSO treated. ‘S’ represents ‘synergy’ as a result of combination treatment evaluation using the Bliss independent model. All results are from three independent experiments.

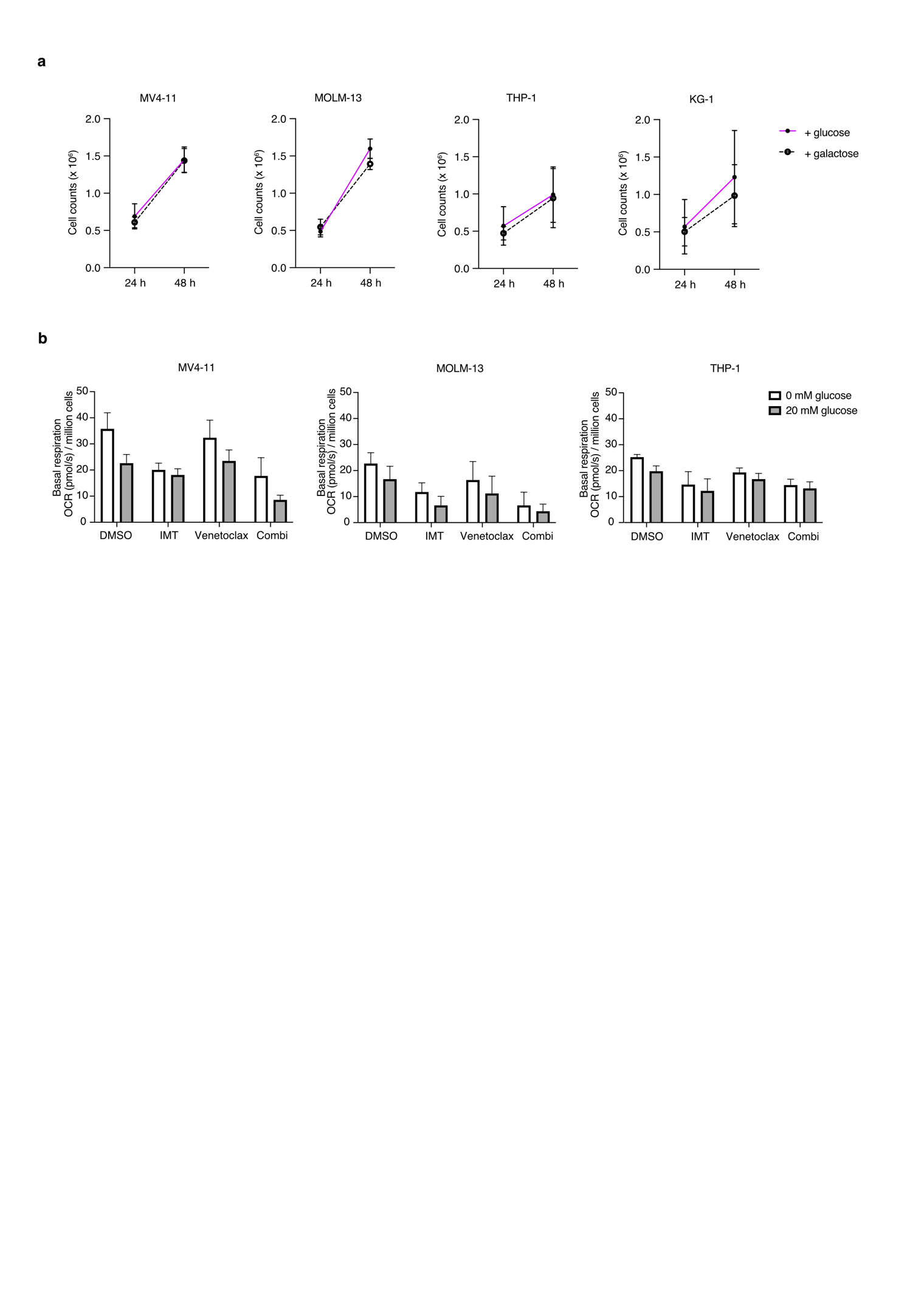

**Figure S3.** (a) AML cell lines were cultured in growth medium supplemented with 5 mM glucose or galactose. Cell counts were performed after 24 and 48 h using NucleoCounter. (b) AML cell lines were treated with either IMT (500 nM), venetoclax (5 nM), or a combination of both for four days. DMSO was added to control wells. Oxygen consumption rate (OCR) was measured at the basal level in the absence/presence of a high concentration of glucose (20 mM) using an Oroboros O2K instrument. All results are from three independent experiments. Statistical tests on all comparisons showed non-significant differences.

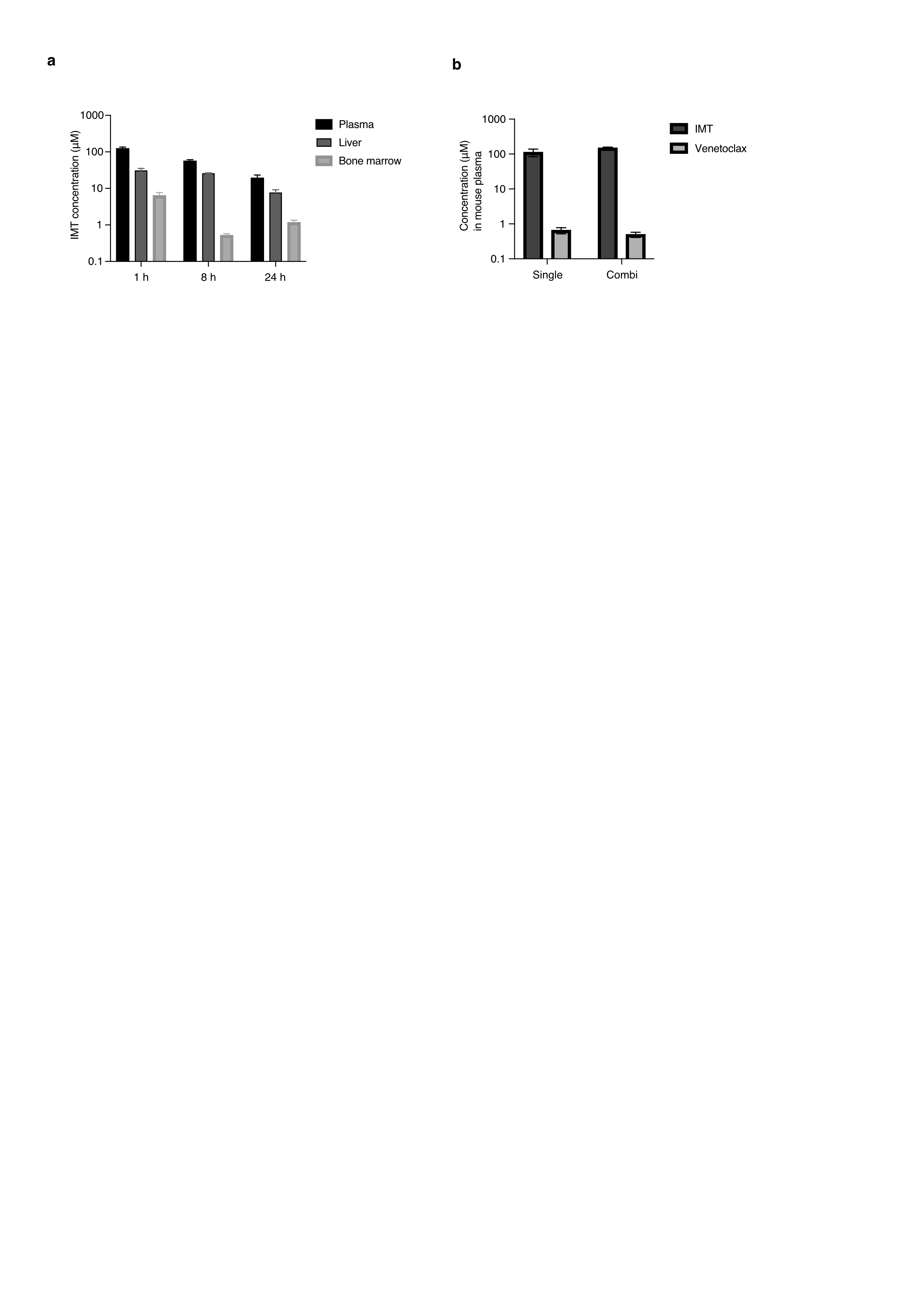

**Figure S4.** (a) *In vivo* pharmacokinetics profile of IMT. Concentration of IMT in plasma, liver, and bone marrow of NSGS mice 1, 8, and 24 h post-administration. (b) Pharmacokinetics of IMT and venetoclax in the plasma of NBSGW mice, 8–11 h post-administration of single compounds or a combination of both compounds. Results are from at least three mice in each group.

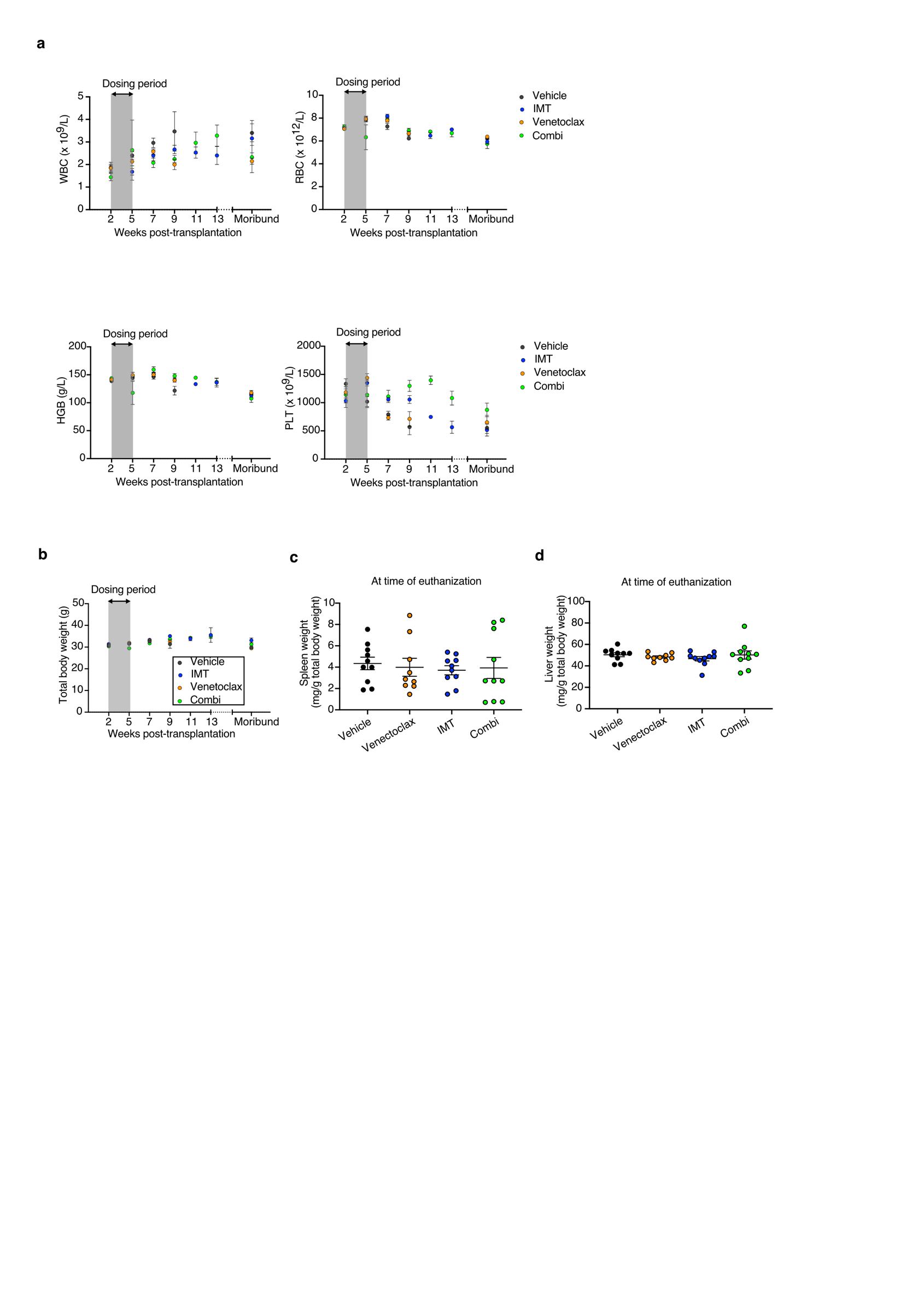

**Figure S5.** Blood values, total weights, and spleen and liver weights of treated CDX mice are not significantly different to those of vehicle-treated mice. (a) Peripheral blood (PB) values of mice shown with white blood count (WBC), red blood count (RBC), hemoglobin (HGB), and platelets (PLT). (b) Total body weights of mice. (c) Ratio of spleen to total body weight. (d) Ratio of liver to total body weight in all treatment groups.

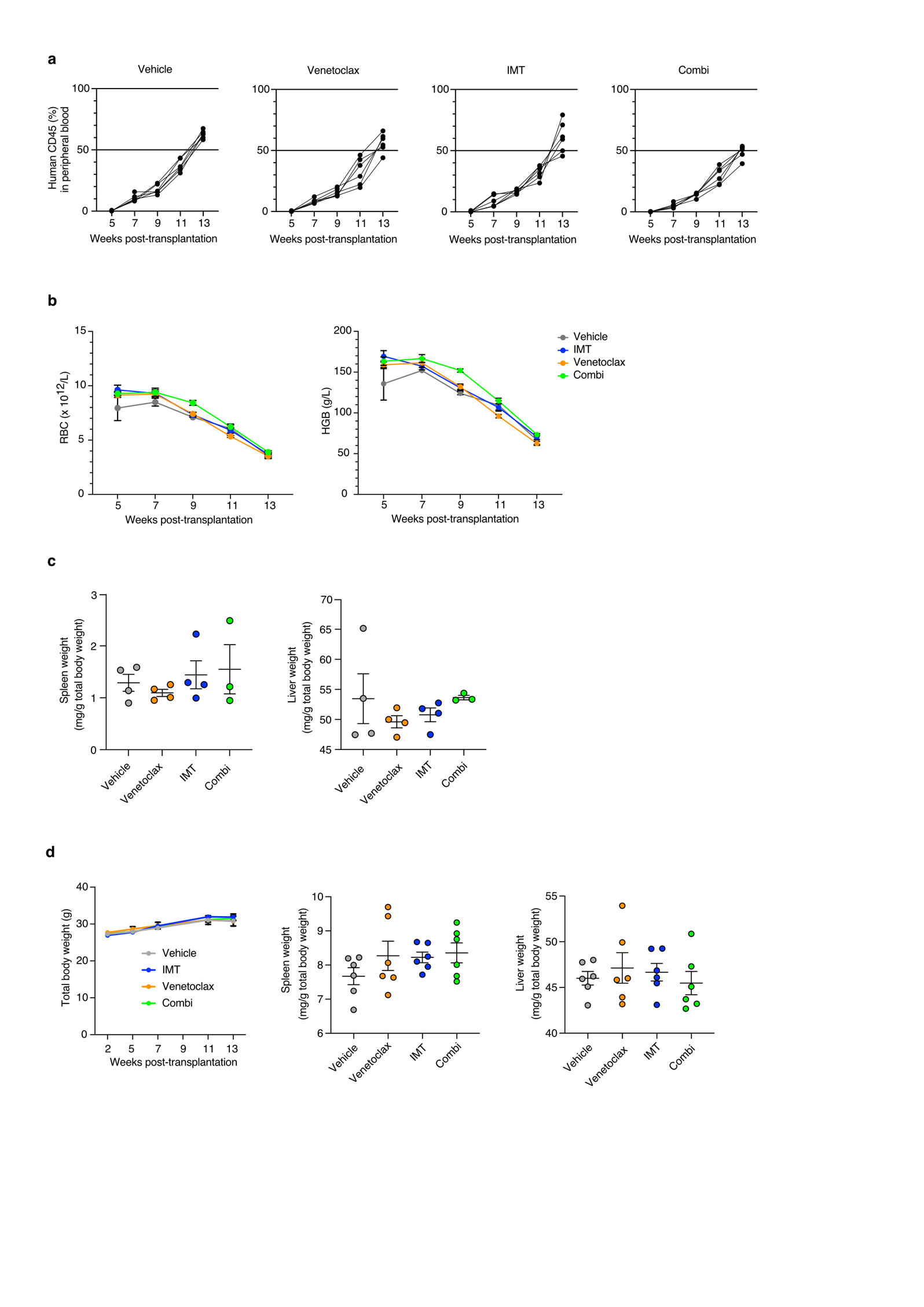

**Figure S6.** (a) Engraftment of hCD45 cells in PB was monitored by flow cytometry at regular intervals until the appearance of disease progression. (b) Red blood counts (RBC) and hemoglobin (HGB) were evaluated until the disease progression. (c) Ratio of spleen and liver weight to total body weight at the end-point. (d) Total body weight, and ratio of spleen and liver to total body weight in the survival group.

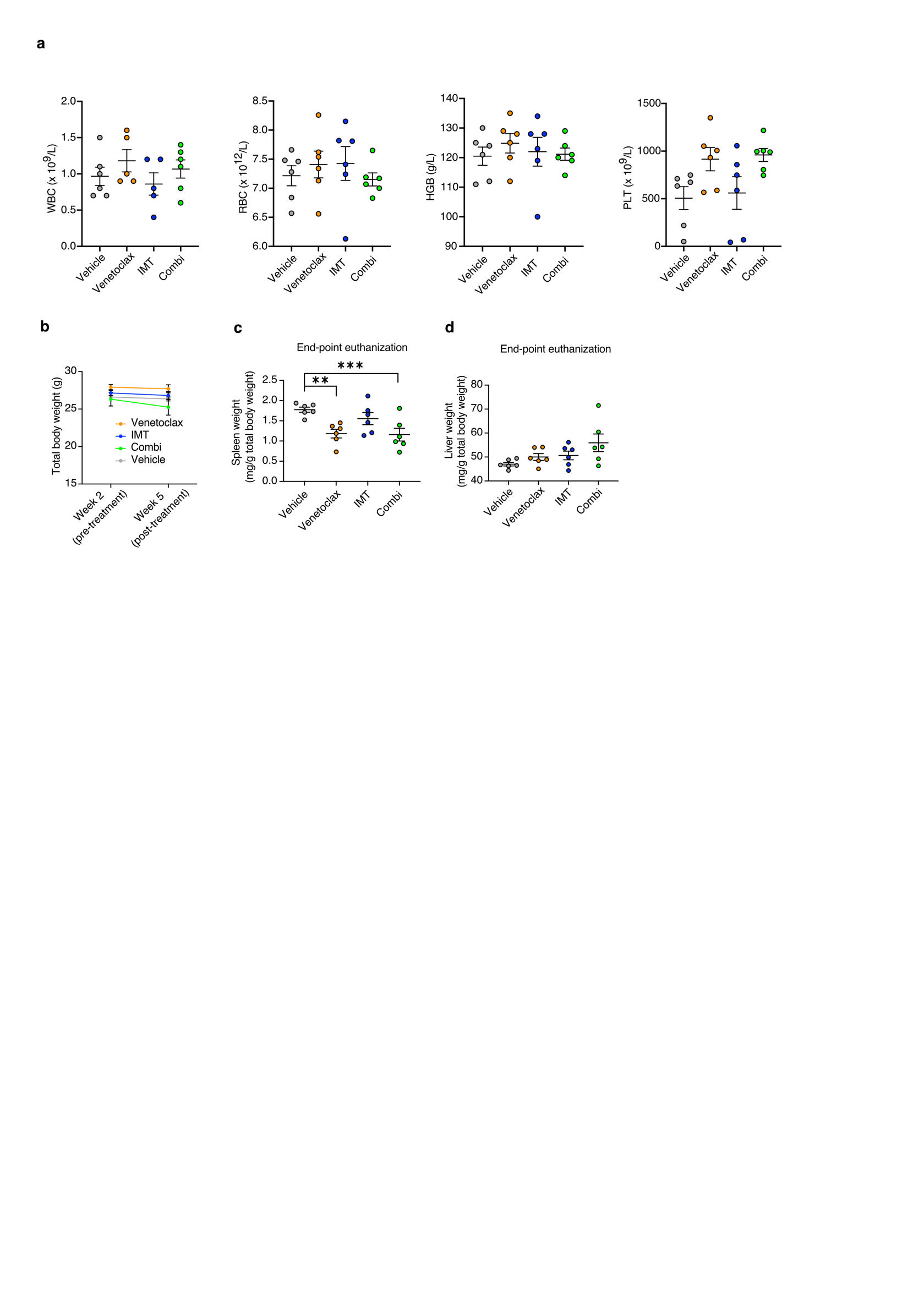

**Figure S7.** (a) Blood values (WBC, RBC, HGB, and PLT), (b) total weights, and (c) spleen and (d) liver weights (mg/g total body weight) evaluated in AML PDX mice after treatment (end-point).

**Table S1**. Summary of activity, ADME (absorption, distribution, metabolism, and excretion) and PK (pharmacokinetics) properties of IMT1B (LDC203974) and LDC204857 (IMT)

|  | IMT1B | LDC204857 |
| --- | --- | --- |
| MW [g/mol] | 474 | 488 |
| SolRank pH 7.4 [µM] | 484 | 484 |
| ClogP | 3,72 | 3,69 |
| PSA [Å²] | 97,1 | 97 |
| biochem hPOLRMT assay IC_50_ | 8.5 nM | 6 nM |
| cellular qRT-PCR assay IC_50_ (mt-7S) | 1,7 nM | 4,6 nM |
| CaCo A2B | 3,6 - 7,6 | 2 - 3,4 |
| CaCo efflux ratio | 2,6 - 6,4 | 3,5 - 7,1 |
| plasma protein binding [%] (human, mouse, rat) | 94(h), 88(m), 90(r) | 98(h), 90(m), 99(r) |
| plasma stability [% at 1h] (h,m,r,d) | 101(m), 98(r) | 100(h), 103(m), 107(r) |
| MS-I *human* [µl/min/mg] metabolic stability phase I | 0,5 | 4,3 |
| MS-I *mouse* [µl/min/mg] metabolic stability phase I | 4,8 - 6,1 | 3,5 |
| MS-I *rat* [µl/min/mg] metabolic stability phase I | 0 | 22 |
| MS-II - glucuronylation  metabolic stability phase II (% remain; h,m,r) | 94(h), 89(m), 88(r) | 96(h), 86(m), 91(r) |
| mouse i.v. (1 mg/kg): CL [l/h/kg] | 0,44 | 0,01 |
| mouse i.v. (1 mg/kg): t1/2 [h] | 1,9 | 12,4 |
| mouse i.v. (1 mg/kg): V_ss_ [l/kg] | 0,86 | 0,13 |
| mouse p.o. (10 mg/kg): t1/2 [h] | 2,7 | 12,4 |
| F [%] p.o. mouse/rat/dog | 100 (m) 25,5 (r) 24,7 (d) | 70,5 (m) 66 (r) 43 (d) |
| CYP 1A2/ 2B6/ 2C8/ 2C9/ 2C19/ 2D6/ 3A4 [µM] | >50 | >50 |
| hERG IC_50_ (µM) | >50 (patch clamp) | >50 (patch clamp) |
| PBMC IC_50_ (µM) | >30 | >30 |

**Table S2.** List of differentially expressed genes in comparison of combination vs vehicle-treated PDX samples.

| Name | log2FoldChange | pvalue | padj | diffexpressed |
| --- | --- | --- | --- | --- |
| SEPTIN3 | -5,2446 | 0,000143 | 0,028895 | down |
| IFI27 | -4,5104 | 0,000185 | 0,034282 | down |
| TRUB2 | -3,7745 | 3,53E-05 | 0,010156 | down |
| CD1E | -3,615 | 0,000141 | 0,02868 | down |
| CLEC10A | -3,4768 | 8,12E-09 | 1,17E-05 | down |
| RGL1 | -3,1916 | 5,29E-05 | 0,014265 | down |
| PARM1 | -3,1624 | 1,33E-06 | 0,00075 | down |
| ZNF22-AS1 | -3,0681 | 0,0001 | 0,022723 | down |
| ALOX5 | -2,9861 | 2,35E-06 | 0,001146 | down |
| PDK4 | -2,7552 | 3,41E-07 | 0,000267 | down |
| F13A1 | -2,6199 | 2,36E-07 | 0,00021 | down |
| S100A9 | -2,4332 | 3,58E-10 | 6,62E-07 | down |
| SIPA1L1 | -2,2632 | 1,86E-07 | 0,000172 | down |
| RAB7B | -2,2543 | 2,11E-05 | 0,006806 | down |
| TRAPPC5 | -2,1821 | 0,000161 | 0,031331 | down |
| TMEM132A | -2,0496 | 2,17E-05 | 0,006814 | down |
| PRLR | -2,0003 | 2,01E-05 | 0,006667 | down |
| MPEG1 | -1,9932 | 1,88E-05 | 0,006335 | down |
| DEPP1 | -1,9642 | 2,15E-09 | 3,47E-06 | down |
| CYBB | -1,9084 | 3,52E-08 | 4,14E-05 | down |
| PHOSPHO1 | -1,8503 | 3,61E-05 | 0,010165 | down |
| RASSF4 | -1,8167 | 1,34E-07 | 0,000128 | down |
| IL10RA | -1,8003 | 6,68E-06 | 0,002662 | down |
| S100A10 | -1,7372 | 0,000106 | 0,023529 | down |
| RARA | -1,7106 | 2,94E-06 | 0,00136 | down |
| FCER1G | -1,6937 | 0,000297 | 0,049974 | down |
| MX1 | -1,6686 | 2,51E-06 | 0,001185 | down |
| SAMHD1 | -1,6579 | 6,54E-12 | 1,88E-08 | down |
| SAMD9L | -1,5997 | 2,47E-05 | 0,007525 | down |
| DTX4 | -1,594 | 2,71E-07 | 0,000233 | down |
| ITGAX | -1,5818 | 0,000129 | 0,02712 | down |
| CST3 | -1,5817 | 1,51E-06 | 0,00083 | down |
| KCTD12 | -1,5703 | 4,45E-07 | 0,000339 | down |
| AHNAK | -1,5683 | 2,03E-17 | 2,63E-13 | down |
| LRP1 | -1,5533 | 0,000205 | 0,037343 | down |
| FCN1 | -1,5469 | 4,42E-06 | 0,001907 | down |
| CPOX | 1,6141 | 0,000267 | 0,045695 | up |
| RCC2-AS1 | 2,2152 | 6,47E-07 | 0,00045 | up |
| PRKRIP1 | 2,2897 | 0,000258 | 0,04489 | up |
| PRORP | 2,82 | 0,000134 | 0,027806 | up |
